# *Thermococci*-to-*Clostridia* Pathway for the Evolution of the Bacteria Domain

**DOI:** 10.1101/2022.10.20.512992

**Authors:** Tze-Fei Wong, Chung-Kwon Chan, Hong Xue

**Affiliations:** Division of Life Science, Hong Kong University of Science and Technology, Clear Water Bay, Hong Kong, China

## Abstract

With the identification of an archaeal Last Universal Common Ancestor phylogenetically related to *Methanopyrus*, the origin of Bacteria becomes a choice between independent emergence versus descent from Archaea. Recently, use of the similarity between paralogous valyl-tRNA synthetase (VARS) and isoleucyl-tRNA synthetase (IARS) as a measure of the ages of bacteria indicated that an Ancestral Bacterial Cluster centred at clostridial *Mahella australiensis* (Mau) were the oldest bacteria. Clostridial *Thermincola potens* (Tpo) also displayed an elevated similarity VARS-IARS bitscore. Overall, the high-bitscore bacteria dominated by *Clostridia* comprised a number of hydrogen producers. A search for archaea capable of hydrogen production that might be ancestral to the Bacteria domain yielded candidates led by *Thermococci* which, like *Clostridia*, form hydrogen through dark fermentation. A two-domain VARS tree based on *Mahella*, *Thermincola*, a broad spectrum of archaea together with both well known and newly reported species of *Thermococci* allocated these two *Clostridia* to a minor-Thermococcal division on the tree. The kinship between *Thermoccoci* and *Clostridia* suggested by this allocation was substantiated by conserved oligopeptide segments on their VARS sequences. It was therefore suggested that a *Thermococci*-to-*Clostridia* evolutionary pathway brought about the emergence of the Bacteria domain.

---

Search for the roots of the three biological domains has been a multi-decade endeavor. According to the proposal that bacteria were the earliest lifeforms, the Last Bacterial Common Ancestor (LBCA) would be equivalent to the Last Universal Common Ancestor (LUCA) of all three domains^1^. A series of bacterial groups have also been suggested as the oldest bacteria, including *Aquifex* and *Thermotoga*^2^; *Planctomycetales*^3^; Gram-positive bacteria^4^; benthic cyanobacteria^5^; a bacterium between Gracilicutes and Terrabacteria near *Fusobacteriota*^6^; and *Megasphaera elsdenii* and *Clostridium tartarivorum* based on gene doubling^7^.

The identification of a LUCA proximal to *Methanopyrus kandleri* (Mka; see Supplementary Tables 1 and 2 for names and abbreviations of microbes), a habitant of marine hydrothermal vents, on the basis of isoacceptor tRNAs^8^ was supported by: the top ranking VARS-IARS bitscore of 473 of Mka among 5,000 species of organisms^9^, which also furnished evidence for the VARS-IARS bitscore as a valuable measure of the relative ages of organisms; usages of the GNN and UNN triplet anticodons by Mka to decode all family boxes of tetracodons, and two neighboring tRNAs in sequence space to translate the well separated UCN and AGY Ser codons on the genetic code^10^; the oldest age of 2.8 Gya of the Mka lineage among archaea^11^; and the tracing of protein families undisturbed by horizontal gene transfer (HGT) back to a LUCA in an environment enriched with hydrogen, carbon dioxide and iron as in a hydrothermal setting^12^. The descent of bacteria from archaea was also consistent with the location of an Ancestral Bacteria Cluster centred at the Clostridial species Mau with a lower VARS-IARS bitscore of 378, closely followed by that of Tpo^9^. Accordingly, the objective of the present study was to search for archaeons that might be candidate Archaeal Progenitors of bacteria with resemblances between their cellular characteristics and those of primitive *Clostridia*. The results obtained showed that the Bacteria domain arose from archaeons phylogenetically located within or near the *Thermococci*.

## Biological attributes of LBCA

The bacteria with the highest VARS-IARS bitscores, headed by Mau and Tpo, were dominated by *Clostridia*. Among non-Firmicute bacteria, only *Thermotogae* appeared on the top-25 list (Table 1). In comparison, *Fusobacteria* attained a bitscore of 312 for *F. ulcerans*; *Cyanobacteria* attained a bitscore of 252 for *Synechococcus sp CC9605*; and *Planctomycetes* attained a bitscore of 201 for *Sedimentisphaera salicampi*. The low bitscore of 148 for *Aquifex aeolicus* was not unexpected in view of its evolution of a substantial oxic network for aerobic metabolism^13^. Although the primitive character of clostridial metabolism among bacteria is clearly regonized^14^, not all *Clostridia* exhibited high VARS-IARS bitscores: the pathogenic *Clostridia* were characterized by more modest bitscores likely on account of the tortuous evolution of their VARS and/or IARS in the course of adapting to a variety of host organisms, in agreement with the phylogenetic tree of *Firmicutes* based on phosphoglycerate kinase where *C. thermocellum* was clustered with *Thermoanaerobacter tencongensis* at a lower branching position than *C. tetani* and *C. perfringens*^15^. Since the G+C content, when estimated from genome sequence, varied by no more than 1% within species, it represented a useful taxonomic descriptor^16^; and the higher G+C contents of 55.5% for Mau and 45.9% for Tpo compared to the 28.6% for *C. tetani*, 28.6% for *C. botulinum* and 27.7-28.7% for *C. perfringens* were consistent with the latter species having to adapt to different hosts. Notably, a number of the top-bitscoring bacteria, such as *Halobacteroides halobius*^17^, *Halothermothrix orenii*^18^, *Carboxydocella thermautotrophica*^19^, *Caldicellulosiruptor*, *Themoanaerobacter* and the non-Firmicute *Thermotogae* are hydrogen-producers^20^. While growth with CO led to the formation of acetate in most *Moorella thermoacetica* strains, such growth produced mainly hydrogen in its atypical AMP strain^21^; and the same-genus *Moorella stamsii* displayed an ability to ferment a range of sugars to yield both hydrogen and acetate^22^. The major end products of pyruvate fermentation by Mau included H_2_, CO_2_ and acetate^23^. Tpo contributed to a hydrogen producing biocathode^24^; and *Thermincola carboxydiphila*, a close relative sharing 99% SSU rRNA identity with Tpo, could oxidize CO with H_2_O to form CO_2_ and H_2_S^25^.

**Table 1.**
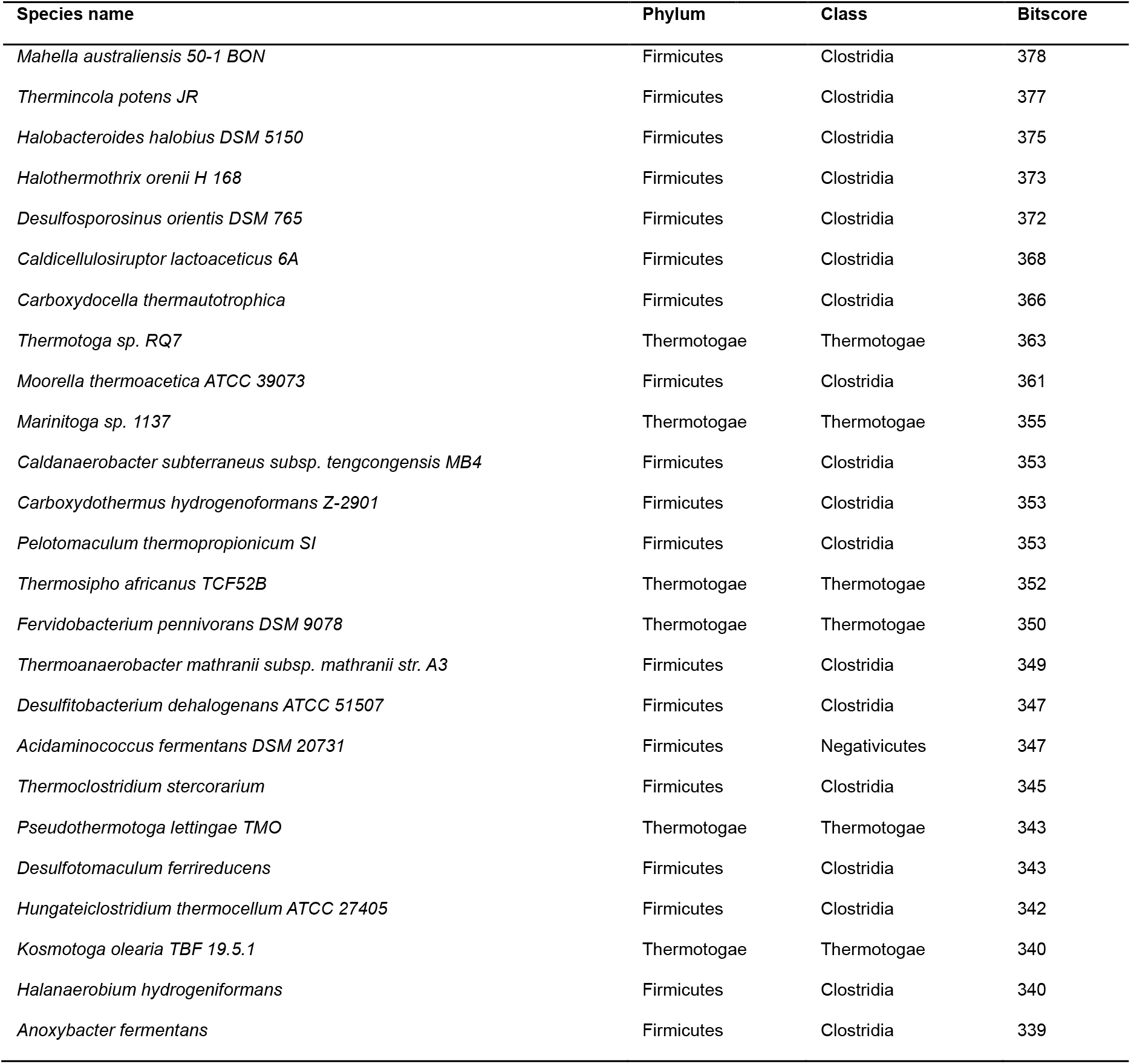
Bacteria with top VARS-IARS bitscores.

## The attributes of Archaeal Progenitor

Since primitive *Clostridia* and plausibly LBCA are anaerobic chemoorganotrophs in possession of a glycolytic sequence and an inclination to produce hydrogen, the Archaeal Progenitor of bacteria might expectedly share some of these cellular attributes. Microbial formation of hydrogen can proceed through biophotolysis as in cyanobacteria, photofermentation as in purple non-sulfur bacteria, electrohydrogenesis involving bacterial consortia and *Proteobacteria* with outer membrane c-type cytochromes, and dark fermentation as in *Clostridia*^26^. Numerous archaeons are hydrogenotrophic methanogens^27^, with the exception of hydrogen-cycling *Methanosarcina* where intracellular production of hydrogen is coupled to its extracellular participation in generating a transmembrane electrochemical gradient^28^. The leading archaea in the development of glycolysis are *Thermococcales* and *Desulfurococcales*, which can degrade 100% of glucose via a modified Embden-Meyerhof-Parnas (EMP) pathway^29^. Since *Desulfurococcales* employ hydrogen to reduce sulfur, whereas *Thermococcales* such as *Pyrococcus furiosis* and *Thermococcus kodakarensis*, generate hydrogen through ferredoxin-NADH based on dark fermentation^26^, *Thermococcales* could be more probable than *Desulfurococcales* as origin of bacterial glycolysis. *Thermococcus* could be more probable than *Pyrococcus* as well in view of the replacement of glyceraldehyde 3-phosphate (GAP) dehydrogenase by GAP: ferredoxin oxidoreductase in the latter’s glycolytic mechanism^20^. Although *Clostridia* utilized an EMP pathway for glycolysis with ATP-dependent kinases, while *Thermococci* preferred to utilize ADP-dependent kinases, the ADP-dependent phosphofructokinase of *Thermococcus kodakarensis* retained 20% activity when ADP was replaced by ATP^30^. Therefore *Thermococci* were faced with relatively few competing candidate Archaeal Progenitors capable of bequeathing to the bacteria both a modified EMP pathway and a dark fermentation mechanism for producing hydrogen. The comparable G+C contents of different *Thermococcus* clades at 45.5%-59.6%^31^, in the vicinity of the G+C contents of Mau and Tpo at 55.5% and 45.9% respectively, would remove any notion that *Firmicutes* are necessarily low in G+C, and facilitate a potential transformation of *Thermococci* as Archaeal Progenitor to form *Clostridia*.

## Importance of high biodiversity sites

Mau was isolated from an oil reservoir^23^, *Clostridium kogasensis* from the soil under a corroded gas pipeline^32^, and *Thermococcus sibiricus* from a high temperature oil reservoir^33^. *Thermococcus* and *Firmicutes* were among the most plentiful archaea and bacteria respectively in reservoirs formed by the pressurized water employed to displace oil from the oil-bearing strata^34^, and *Thermincola* were abundant in reservoirs above 90°C^35^. Thus geobiological sites favorable for a transition from *Thermococci* to *Clostridia* could include niches bearing some petroleum-bearing niches. Notably, a survey of fifteen *Thermococcus* species revealed that none of the species originating outside of hydrothermal vents metabolized maltose as energy substrate, while a majority of the species originating from hydrothermal vents, including *T. aggregans*, *T. guaymasensis*, *T. fumicolans*, *T. hydrothermalis* and *T. profundus* all metabolized maltose^36^, suggesting that high biodiversity sites such as the vents would enhance divergence of the chemoorganotrophic *Thermococcus* from their cellular norm to develop breakthrough properties including bacterial-type sugar glycolysis based on the EMP pathway.

In this regard, the marine hydrothermal vents at Guaymas Basin are known to release an abundance of CO_2_, H_2_ and low molecular-weight peptides, hydrocarbons and carbohydrates when magmatic sills intruded into organic-rich sediments up to 28 to 7 kyr ago^37,38^, and these compounds gave rise to elevated biodiversity^39,40^. In a single study, five novel *Thermococcus* species were identified from these vents based on SSU rRNA, elongation factors EF-1alpha and EF-2, which indicated a rapid rate of *Thermococus* evolution^41^. More recently, based on 37 single-copy genes including the VARS and IARS genes, a series of metagenome-assembled genomes (MAGs) of *Thermococci* and *Euryarchaeota* from these vents were sequenced^42,43^. The terrestrial subsurface has been revealed likewise as a hotspot of anaerobic biodiversity, where a single site was known to harbor much of the tree of life^44^: the cold water CO_2_-driven Crystal Geyser in Utah allowed the sampling of 104 different phylum-level lineages of archaea and bacteria from different depths of the subsurface in the form of genome-resolved metagenomes and single amplified genomes (SAGs), including a novel *Euryarchaeota archaeon* genome with a bitscore of 403 (viz. bit403)^45,46^.

## VARS tree and sequence alignment

Archaeal and bacterial species that produce hydrogen through dark fermentation are relatively few. To examine whether the resemblance between *Thermococci* and primitive *Clostridia* represented convergent evolution or phylogenetic relatedness, a two-domain maximum parsimony VARS tree was built using the LUCA-proximal *Methanopyrus kandleri* as root. There were a major-Thermococcal division consisting of familiar and novel species of *Thermococci* and *Euyarchaeota* such as *T. litoralis* and *T. sibiricus*, and a minor-Thermococcal division consisting of novel species such as *T. archaeon* B48 G16 from Guaymas Basin and *Euryarchaeota archaeon* bit403 from Crystal Geyser. Different archaeal species with 300 or higher VARS-IARS bitscores were included preferentially in the tree in order to minimize the impact of horizontal gene transfers (HGTs); since HGTs involving both the VARS and IARS genes would be unlikely, while HGTs involving either VARS or IARS alone would lower the similarity bitscore between VARS and IARS substantially, same-species VARS-IARS pairs that have managed to maintain a high bitscore could have been protected relatively against HGT perturbation.

In confirmation that the allocation of Mau and Tpo to the minor-Thermococcal division on the VARS tree was guided by kinship instead of HGT, Figure 2 shows oligopeptide segments I-V excerpted from the full length VARS alignment in Supplementary Figure 1:

*Oligo I*. Eight of the amino acid positions in each of the *Mahella* and *Thermincola* nanopeptides were identical to those of *Thermococcus sibiricus*.
*Oligo II*. The third position was ‘V’ in the major-Thermococcal division, but ‘V’ or ‘C’ in the minor-Thermococcal division, which supported the allocations of *Mahella* and *Thermincola* into the minor-Thermococcal division.
*Oligo III*. The decapeptides on *Mahella* and *Thermincola* VARSs were identical to those on *Paleococcus pacificus* and *T. archaeon* B89 G9.
*Oligo IV*. The heptapeptide on *Mahella* was identical to those on *T. sibiricus*, *T. archaeon* B45 G15, *T. sp. 2319×1*, *T. litoralis*, T. archaeon bit391, *T. kodakarensis* and *P. furiosis*.
*Oligo V*. The heptapeptide on *Mahella* differed at only one position from those on *T. sp. 2319×1*, *T. litoralis*, *T. archaeon* bit391, *P. pacificus, T. archaeon* B89 G9, *T. kodakarensis* and *P. furiosis*.

Thus Oligos I-V on *Thermococci* VARS resembled their counterparts on the primitive *Clostridia* genera *Mahella* and *Thermincola*, especially *Mahella*, even more than their counterparts on the *Crenarchaeota* species *Thermofilum pendens*, *Themoprotei archaeon* bit383 and *Crenarchaeota archaeon* bit341. Notably, while HGTs occur most frequently between closely related species^47^, the *Mahella* sequences in Oligos I-V showed similarities toward not only the *Thermococci* sequences but also most of the other archaeal sequences as well, albeit to a lesser extent. Therefore, the probability of Oligos I-V arising from cross-domain HGTs between *Clostridia* and the wide range of archaea would be vanishingly small, establishing thereby the phylogenetic proximity of *Mahella* to a *Thermococci*-related LBCA^48^.

## Discussion

Few archaeon-bacterium pairs are so comparable in their biologies or genomes that they would be suggestive of an evolutionary precursor-product relationship, and this was the case even with pairings between *Thermococcus* and the pathogenic *Clostridium* species. In contrast, the phylogenetic proximity between *Thermococcus* and primitive *Clostridia* was supported by: the top VARS-IARS similarity bitscores of the latter among bacteria^9^; the antiquity of the clostridial metabolic network^*14*^; their shared anaerobiosis and thermophilicity (*Thermococci* can grow under hyperthermophilic or thermophilic conditions, while Mau is a moderate thermophile with an optimal growth temperature of 50°C, and Tpo was isolated from a thermophilic microbial fuel cell); their medium-level G+C content; their adaptability to petroleum containing sites; the presence of a modified EMB pathway in *Thermococci* with the same intermediates as bacterial glycolysis; and their hydrogen production through ferredoxin-mediated dark fermentation. With such an extensive match between their cellular profiles, their co-location on the two-domain VARS tree (Figure 1) and striking resemblance in Oligos I-V (Figure 2) were by no means a surprise.

**Figure 1.**
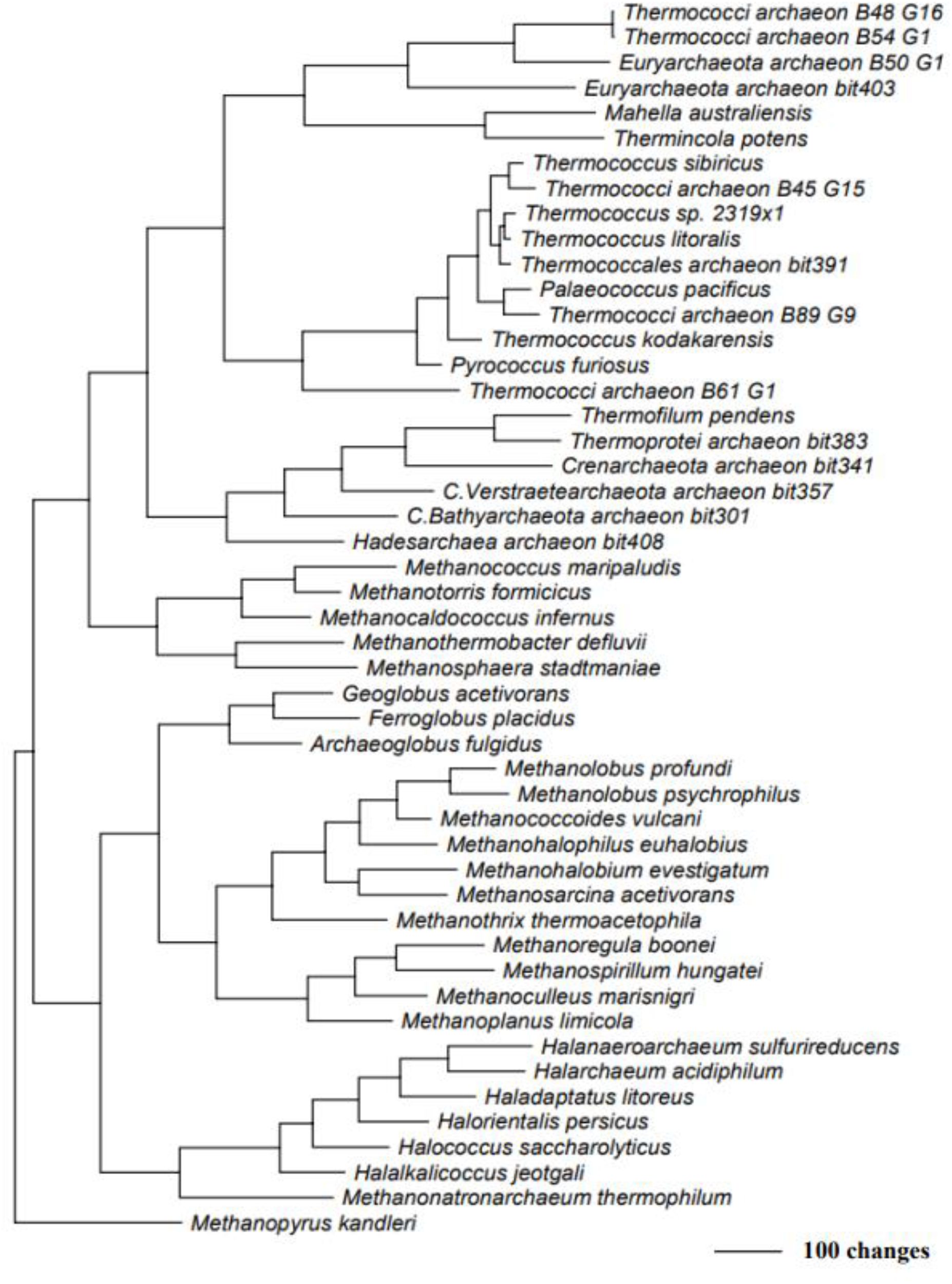
Two-domain VARS tree based of the VARS sequences of a wide range of archaea together with the Clostridial Mau and Tpo, and *Methanopyrus kandleri* as root. VARS sequences and bitscores are shown in Supplementary Tables 1 and 2.

**Figure 2.**
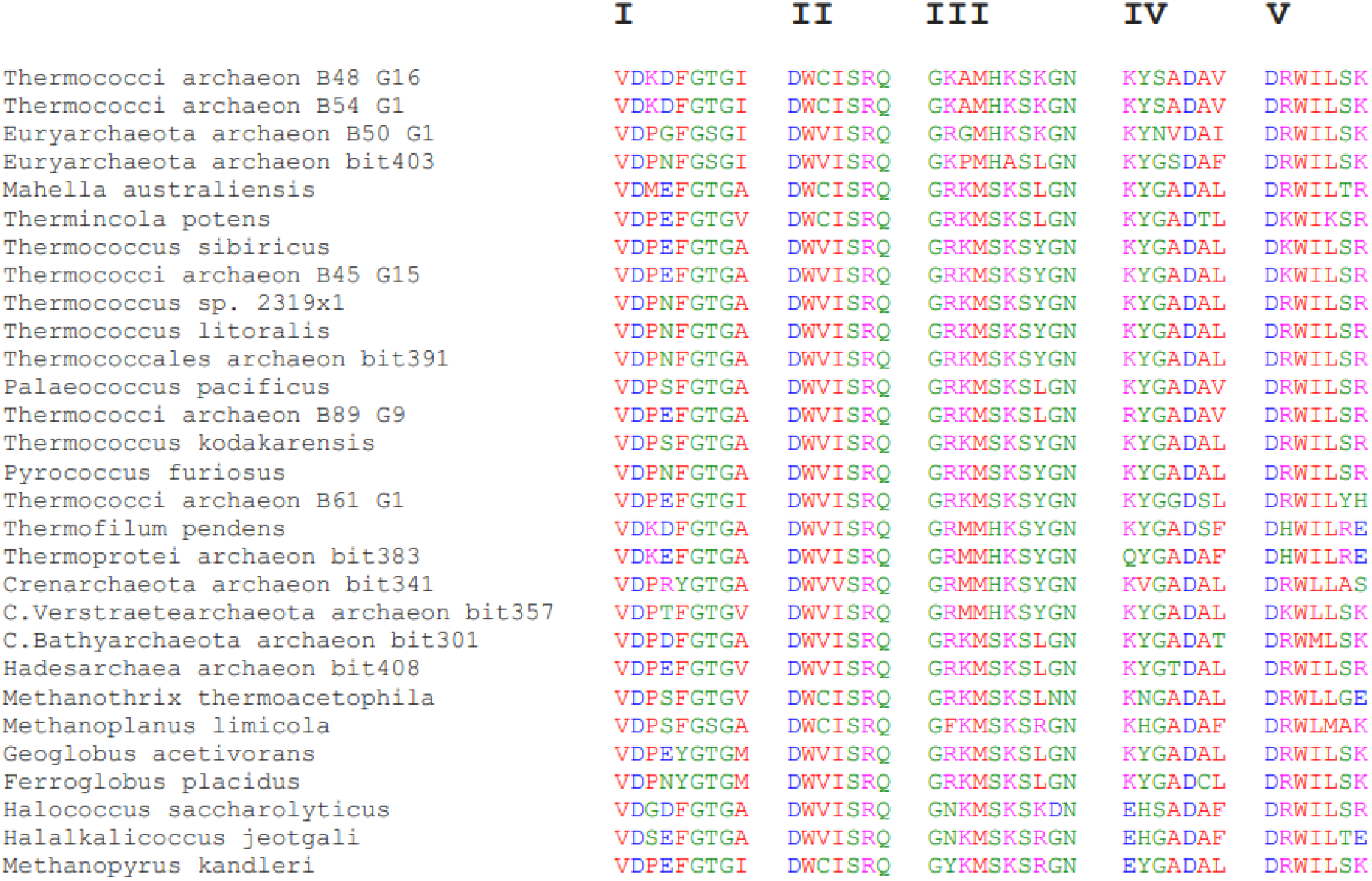
Five aligned Oligopeptide segments from some microbes on the VARS tree were excerpted from the full length VARS alignments in Supplementary Figure 1, starting at positions 292, 433, 575, 594 and 664 respectively.

Accordingly, the sister clades of *Thermococci* and primitive *Clostridia* on the VARS tree (Figure 1), and the striking conservation of Oligos I-V between them (Figure 2) have provided strong evidence for the emergence of Bacteria from Archaea via a *Thermococci*-to-*Clostridia* evolutionary pathway, as well as innovative *Thermococci* evolution at high biodiversity sites such as the Guaymas Basin and the terrestrial subsurface. Such a pathway will contain by necessity many deep divergences between the Archaea and Bacteria domains that need to be explained in terms of the genomic changes between them. Clearly, wide-ranging investigation including the continued search for more *Thermococci-* and *Euryarcheota-*like *Clostridia*, and more *Clostridia-*like *Thermococci* and *Euryarcheota*, will be required to decipher how the extraordinarily complex transformation of an archaeal genome into a bacterial genome came to be accomplished.

## Methods

VARS sequence alignment was performed with CLUSTALX. A maximum parsimony analysis was conducted using PAUP* version 4.0a169 win32^48^. Heuristic searches with 10 random taxon addition replicates and tree bisection-reconnection swapping were applied.

## Acknowledgements

We thank Ms Si Chen for server management. This research was funded by grant number 5112-703-0110D-42002 of the University Grants Council of Hong Kong to Applied Genomics Center of Hong Kong University of Science and Technology.

## Author contributions

Conceptualization, T.F.W. and H.X.; data analysis, C.K.L.; writing, T.F.W., C.K.L. and H.X. All authors have read and agreed to the published version of the manuscript.

## Additional Information

All data generated or analysed during this study are included in this published article and its supplementary information files. The following are available online. Supplementary Figure 1: Full length alignment of VARSs from various organisms. The numbering of amino acid residues is based on the VARS of Crenarchaeota archaeon bit341. Supplementary Table 1: Bacterial species with relatively high VARS-IARS bitscores (ref. 9). Supplementary Table 2: Archaeal species with relatively high VARS-IARS bitscores. See ref. 43 for sequences of *C.Hydrothermarchaeota archaeon_B51_G15*, *Thermococci archaeon_B45_G15, Thermococci archaeon_B89_G9, Thermococci archaeon_B61_G1*, *Thermococci archaeon_B88_G9*, *Thermococci archaeon_B48_G16*, *Thermococci archaeon_B54_G1* and *Thermoprotei archaeon_B75_G16*; ref. 46 for sequence of *Euryarchaeota archaeon_bit403*; and ref. 9 for remaining entries.

## References

1. Woese, C. R., Kandler, O. & Wheelis, M. L. Towards a natural system of organisms: proposal for the domains Archaea, Bacteria and Eucarya. Proc. Natl Acad. Sci. USA 87, 4576–79 (1990).

2. Bocchetta, M., Gribaldo, S., Sanangelantoni, A. & Cammarano, P. Phylogenetic depth of the bacterial genera Aquifex and Thermotoga inferred from analysis of ribosomal protein, elongation factor and RNA polymerase subunit sequences. J Mol Evol 50, 366–380 (2000).

3. Brochier, C. & Philippe, H. A non-hyperthermophilic ancestor for Bacteria. Nature 417, 244 (2002).

4. Ciccarelli, F. D. et al. Toward automatic reconstruction of a highly resolved tree of life. Science 311, 1283–87 (2006).

5. Noffke, N., Beukes, N., Bower, D., Hazen, R. M. & Swift D. J. P. An actualistic perspective into Archean worlds-(cyano-) bacterially induced sedimentary structures in the siliciclastic Nhlazatse section, 2.9 Ga Pongola Supergroup, South Africa. Geobiol 6(1), 5–20 (2008).

6. Coleman, G. A. et al. A rooted phylogeny resolves early bacterial evolution. Science 372, eabe0511 Doi: 10.1126/science.abe0511 (2021).

7. Schwartz, R. M. & Dayhoff, M. O. Origins of prokaryotes, eukaryotes, mitochondria and chloroplasts. Science 199, 395–404 (1978).

8. Xue, H., Tong, K. L., Marck, C., Grosjean, H. & Wong, J. T. F. Transfer RNA paralogs: evidence for genetic code-amino acid biosynthesis coevolution and an archaeal root of life. Gene 310, 59–66 (2003).

9. Long, X., Xue, H. & Wong, J. T. F. Descent of Bacteria and Eukarya from an archaeal root of life. Evolutionary Bioinformatics 16, 1–11 (2020).

10. Wong, J. T., Ng, S. K., Mat, W. K., Hu, T., & Xue, H. Coevolution Theory of the Genetic Code at Age Forty: Pathway to Translation and Synthetic Life. Life 6(1), 12–(2016).

11. Blank, C. E. Low rates of lateral gene transfer among metabolic genes define the evolving biogeochemical niches of archaea through deep time. Archaea 2012, article 843539 Doi.org/10.1155/2012/843539 (2012).

12. Weiss, M. C., Preiner, M., Xavier, J. C., Zimorski, V. & Martin, W. F. The last universal common ancestor between ancient Earth chemistry and the onset of genetics. PLOS Genetics 14(8), e1007518 (2018).

13. Raymond, J. & Segrè, D. The effect of oxygen on biochemical networks and the evolution of complex life. Science 311, 1764–67 (2006).

14. Xavier, J. C. et al. The metabolic network of the last bacterial common ancestor. Commun Biol 4, 413 (2021).

15. Wolf, M., Muller, T., Dandekar, T. & Pollack, J. D. Phylogeny of Firmicutes with special reference to Mycoplasma (Mollicutes) as inferred from phosphoglycerate kinase amino acid sequence data. Int J Syst Evol Microbiol 54(3), 871–875 (2004).

16. Meier-Kolthoff, J. P., Klenk, H. P. & Göker, M. Taxonomic use of DNA G+C content and DNA-DNA hybridization in the genome age. Int J Syst Evol Microbiol 64(Pt 2), 352–356 (2014).

17. Oren, A., Weisburg, W. G., Kessel, M. & Woese, C. R. Halobacteroides halobius gen. nov., sp. nov., a moderately halophilic anaerobic bacterium from the bottom sediments of the Dead Sea. Syst Appl Microbiol 5, 58–69 (1984).

18. Cayol, J. L. et al. Isolation and characterization of Halothermothrix orenii gen. nov., sp. nov., a halophilic, thermophilic, fermentative, strictly anaerobic bacterium. Int J Syst Bacteriol 44(3), 534–540 (1994).

19. Toshchakov, S. V. et al. Genomic insights into energy metabolism of *Carboxydocella thermautotrophica* coupling hydrogenogenic CO oxidation with the reduction of Fe(III) minerals. Front Microbiol doi.org/10.3389/fmicb.2018.01759 (2018).

20. Verhaart, M. R. A., Bielen, A. A. M., van der Oost J., Stams, A. J. M. & Kengen, S. W. M. Hydrogen production by hyperthermophilic and extremely thermophilic bacteria and archaea: mechanisms for reductant disposal. Environ Technol 31(8-9), 993–1003 (2010).

21. Jiang, B. et al. Atypical one carbon metabolism of an acetogenic and hydrogenic *Moorella thermoacetica* strain. Arch Microbiol 191, 123–131 (2009).

22. Alves, J. I., van Gelder, A. H., H., Alves, M. M., Sousa, D. Z. & Plugge, C. M. Moorella stamsii sp. nov., a new anaerobic thermophilic hydrogenic carboxydotroph isolated from digester sludge. Int J Syst Evol Microbiol 63(11), 4072–76 (2013).

23. Bonilla Salinas, M. et al. Mahella australiensis gen nov., sp. nov., a moderately thermophilic anaerobic bacterium isolated from an Australian oil well. Int J Syst Evol Microbiol 54(6), 2169–73 (2004).

24. Fu, Q. et al, Bioelectrochemical analyses of a thermophilic biocathode catalyzing sustainable hydrogen production. Int J Hydrogen Energy 38(35), 15638–45 (2013).

25. Sokolova, T. G. et al. Thermincola carboxydiphila gen. nov., sp. nov., a novel anaerobic carboxydotrophic hydrogenic bacterium from a hot spring of the Lake Baikal area. Int J Syst Evol Microbiol 55(5), Doi10.1099/ijs.0.63299-0 (2005).

26. van Niel E. W. J. Biological processes for hydrogen production. Adv Biochem Eng Biotech DOI: 10.1007/10_2016_11 (2016).

27. Richards M. A. et al. Exploring hydrogenotrophic methanogenesis: a genome scale metabolic reconstruction of Methanococcus maripaludis. J Bacteriol 198: 3379–90.

28. Kulkani, G., Mand, T. D. & Metcalf, W. W. Energy conservation via hydrogen cycling in the methanogenic archaeon Methanosarcina barkeri. mBio 9, e01256.

29. Bräsen, C., Esser, D., Rauch, B. & Siebers, B. Carbohydrate metabolism in Archaea: current insights into unusual enzymes and pathways and their regulation. Microbiol Mol Bio Rev 78(1), 89–175 (2014).

30. Shakir, N. A., Bibi, T., Aslam, M. & Rashid, N. Biochemical characterization of a highly active ADP-dependent phosphofructokinase from Thermococcus kodakarensis. J Biosci Bioeng 129(1), 6–15 (2020).

31. Price, M. T., Fullerton, H. & Moyer, C. L. Biogeography and evolution of Thermococcus isolates from hydrothermal vent systems of the Pacific. Front Microbiol 24, doi.org/103389/fmic.b.2015.00968 (2015).

32. Shin, Y. et al. Clostridium kogasensis sp. nov., a novel member of the genus Clostridium isolated from the soil under a corroded gas pipeline. Anaerobe 39, 14–18 (2016).

33. Miroshnichenko, M. L. et al. Isolation and characterization of Thermococcus sibiricus sp. nov. from a Western Siberia high-temperature oil reservoir. Extremophiles 5(2), 85–91 (2001).

34. Wang, L. Y. et al. Molecular analysis of the microbial community structures in water-flooding petroleum reservoirs with different temperatures. Biogeosci 9(11), 4645–59 (2012).

35. Lin, J. et al. A study on the microbial community structure in oil reservoir developed by water flooding. J Petroleum Sci Engineering 122, 354–359 (2014).

36. Kobayashi, T. Genus I. Thermococcus. In Bergey’s Manual of Systematic Bacteriology 2^nd^ ed., Vol 1. The archaea and deeply branching and phototrophic bacteria, eds. Boone, D., Castenholz, R. W. & Garrity, G. M. Springer pp 342–346 (2001).

37. Einsele, G. et al. Intrusion of basaltic sills into highly porous sediments, and resulting hydrothermal activity. Nature 283, 441–445 (1980).

38. Simoneit, B. R. T. & Lonsdale, P. F. Hydrothermal petroleum in mineralized mounds at the seabed of Guaymas Basin. Nature 295, 198–202 (1982).

39. McKay, L. et al. Thermal and geochemical influences on microbial biogeography in the hydrothermal sediments of Guaymas Basin, Gulf of California. Environ Microbial Rep 8, 150–161 (2016).

40. Lin, Y. S. et al. Near surface heating of young rift sediment causes mass production and discharge of reactive dissolved organic matter. Sci Rep 7, 44864 doi.org/10.1038/srep44864 (2017).

41. Liu, L., Wang, F., Xu, J. & Xiao, X. Molecular diversity of Thermococcales isolated from Guaymas Basin hydrothermal vents. Acta Oceanologica Sinica 32, 75–81 (2013).

42. Dombrowski, N., Teske, A. P. & Baker, B. J. Expansive microbial metabolic versatility and biodiversity in dynamic Guaymas Basin hydrothermal sediments. Nature Comm 9, 4999 (2018).

43. Dombrowski, N., Seitz, K. W., Teske, A. P. & Baker, B. J. Genomic insights into potential interdependencies in microbial hydrocarbon and nutrient cycling in hydrothermal sediments. NCBI Bioproject PRJNA362212 (2017).

44. Anantharaman, K. et al. Thousands of microbial genomes shed light on interconnected biogeochemical processes in an aquifer system. Nature Comm 17, 13219 (2016).

45. Probst, A. J. et al. Differential depth distribution of microbial function and putative symbionts through sediment-hosted aquifers in the deep terrestrial subsurface. Nature Microbiol 3, 328–336 (2018).

46. Probst, A. J. et al. Differential depth distribution of microbial function and putative symbionts through sediment-hosted aquifers in the deep terrestrial subsurface. NCBI Bioproject (2018). PRJNA362739 Euryarchaeota archaeon CG_4_9_14_3_um_filter_38_12(groundwater metagenome).

47. Soucy S. M., Huang, J. & Gogarten, J. P. Horizontal gene transfer: building the web of life. Nature Rev genet 16, 472–482 (2015).

## References

48. Swoford DL. PAUP*: Phylogenetic analysis using parsimony (* and other methods), version 4.0b10 win32. Sunderland. MA:Sinauer Associates (2002).

