## Supplementary Figure 1 for "*Thermococci*-to-*Clostridia* Pathway for the Evolution of the Bacteria Domain"

CLUSTAL O(1.2.4) multiple sequence alignment

```
                                                      10        20        30        40        50        60
                                              .........|.........|.........|.........|.........|.........|
Thermococcus sp. 2319x1                       -------MLPKTYDPNEIEPKWQKFWLEEKIYKYELD--EKRPAYSIDTPPPFTSGTLHL
Thermococcus litoralis                        -------MLPKTYDPNEIEPKWQKFWLEEKIYKYELD--EKRPAYSIDTPPPFTSGTLHL
Thermococcales archaeon bit391                -------MLPKTYDPNEIEPKWQKFWLDEKIYKYELD--EKRPAYSIDTPPPFTSGTLHL
Thermococcus sibiricus                        -------MLSKSYDPNEIESKWQKFWLEEGIYKYRLD--EKRPAYSIDTPPPFTSGTLHL
Thermococci archaeon B45 G15                  -------MLSKTYDPNEIEPKWQKFWLEEEIYKYKLD--EKKPAYSIDTPPPFTSGTLHL
Palaeococcus pacificus                        --------MQKTYNPKDIEPKWQKFWLNEKIYKYELD--EKRPAYAIDTPPPFTSGTLHL
Thermococci archaeonB89 G9                    --------MQKTYDPKEIEPKWQKFWLDERIYKYELD--EKRPPYSIDTPPPFTSGTLHL
Thermococcus kodakarensis                     -------MLPKNYDPNEIEPKWQKFWLDEKIYKYELD--EKKPSYAIDTPPPFTSGTLHL
Pyrococcus furiosus                           -------MLPKNYDPNEIEPKWQKYWLEEKIYKYKLE--PDKPSYAIDTPPPFTSGTLHL
Thermococci archaeon B61 G1                   --------MDKEYNHKEVEKKWIEKWSKSDIYSFKND---GREVFSIDTPPPYTSGVAHM
Thermococci archaeon B48 G16                  --------MLGEYNPKEIEEKVQRFWEEKEIYKFDPD--SEKPVFSIDTPPPTFSGEIHI
Thermococci archaeon B54 G1                   --------MLGEYNPKEIEEKVQRFWEEKEIYKFDPD--SEKPVFSIDTPPPTFSGEIHI
Euryarchaeota archaeon bit403                 ----------MVYEQKSVEQKWQKKWQKMSLYSFVPD--SDKPVYSIDNPPRYASGALHV
Mahella australiensis                         -----MEQLSKAYDPSKVEDKWYTEWMDKGYFRAEVD--KNKQPFCIVMPPPNITGQLHI
Thermincola potens                            --MSAEKNLPTTYNPGEVEKKWYKFWEDNGFFHAEVN--PEKKPFAIVMPPPNVTGALHM
Thermofilum pendens                           --MTVEFRLPKEYNFKAVEGKWQRFWEEKGIYRFDRK-DRSRPVYVIDTPPPYPSGDLHV
Thermoprotei archaeon bit383                  -----MEEFPKRYNFREIEKKWQKTWEEWGLYRFDRE-DKSRPTYSIDTPPPYPSGEVHV
Crenarchaeota archaeon bit341                 MGTGRNYALDKNFDPAAEESKWREYWREHDIYHFDPL-DKSRKTYSIDTPPPYPSGDFHV
C.Verstraetearchaeota archaeon bit357         ------MILPRDYDFRKVEVKWQRKWEEWSIYSFDWE-DFEKPTYAIDTPPPYPSGEFHM
C.Bathyarchaeota archaeon bit301              -----MRELPKDFDILSIEEKWQKKWMELEVYRFHRE-DKEKPPFVIDTPPPYPSGDFHM
Hadesarchaea archaeon bit408                  ---MTKVEVSKDYDPLKVEPRWQKRWEELDLYRFDRE-DKTKPTYVIDTPPPYPSGEFHM
Methanothrix thermoacetophila                 -----MSEISKEYNFKEVEEKWIERWDPSVYY-F--DWGSEKPQYIIDTPPPYPTGNFHI
Methanoplanus limicola                        --MSDTGEIPKNYDPLEVEERWLGIWKEENYY-F--DRNSSKPRFIIDTPPPYPTGNFHI
Geoglobus acetivorans                         --------MEKEYNAHKVEEKWVEQWKDEMYY-F--DWNSEKPHFIIDTPPPYPTGTFHI
Ferroglobus placidus                          --------MEKEYKAREVEDKWLRLWKDDMYY-F--DWDSKKPHFIIDTPPPYPTGSFHI
Archaeoglobus fulgidus                        ------MEIRKDYDAHEVEEKWLKLWKDEMYY-F--DWNSEKPHYIIDTPPPYPTGSFHI
Halococcus saccharolyticus                    -----MTELSETYDPEALEAKWREEWHESDLYRYDDDNDDTDTQYVIDTPPPYPTGDFHI
Halalkalicoccus jeotgali                      --------MDDSYDPGAIEPRWRERWADEDVYRYDGDE--KRPEYVIDTPPPYPTGNLHI
Methanopyrus kandleri                         ---MPGEAPVEDYDPKEIEPKWRERWLEERKYRFEGG--EDRPAFVIDTPPPYPTGELHM
                                                          :.    * :    *     :            : *  **   :*  *:
```

```
                                                  70        80        90        100       110       120
                                          .........|.........|.........|.........|.........|.........|
Thermococcus sp. 2319x1                   GHVLSHTWIDIVARYKRMGGYNVLFPQGFDNHGLPTELKVEKE---FGISKD--QPEEFL
Thermococcus litoralis                    GHVLSHTWIDIVARYKRMGGYNVLFPQGFDNHGLPTELKVEKE---FGISKD--QPEEFL
Thermococcales archaeon bit391            GHVLSHTWIDIVARYKRMRGYNVLFPQGFDNHGLPTELKVEKE---FGISKD--QPEEFL
Thermococcus sibiricus                    GHVLSHTWIDIVARYKRMRGYNVLFPQGFDNHGLPTELKVEKE---FGISKD--QPEEFL
Thermococci archaeon B45 G15              GHVLSHTWIDIVARYKRMRGYNVLFPQGFDNHGLPTELKVEKE---FGISKD--QPEEFL
Palaeococcus pacificus                    GHVLSHTWIDIIARYKRMRGYNVLFPQGFDNHGLPTELKVEKE---FGISKD--EPEKFL
Thermococci archaeonB89 G9                GHVLSHTWIDIVARYKRMRGYNVLFPQGFDNHGLPTELKVEKE---FGISKD--EPERFL
Thermococcus kodakarensis                 GHVLSHTWIDIIARYKRMTGYNVLFPQGFDNHGLPTELKVEKE---FGISKD--QPEKFL
Pyrococcus furiosus                       GHVLSHTWIDIIARYKRMRGYNVLFPQGFDNHGLPTELKVEKE---FGISKD--QPEEFL
Thermococci archaeon B61 G1               GTILGYVWFDMIARFKRMRGFDVYFPHGWDTQGLPTELKVEKE---LGKSAK-KDRSKFR
Thermococci archaeon B48 G16              GHAMSYSQAEFMARYKRMRGYNIFYPMGFDDNGLPTERLVEQK---YKVDIKEIGREKFV
Thermococci archaeon B54 G1               GHAMSYSQAEFMARYKRMRGYNIFYPMGFDDNGLPTERLVEQK---YKVDIKEIGREKFV
Euryarchaeota archaeon bit403             GHAVHYTHIDFAARYRRMKGFNVFFPLCFDCNGIPIEERVEKK---LGITRKDIDRHKFI
Mahella australiensis                     GHAVNNTLQDILIRWRRMQGYNALWIPGTDHASIATEAKIKDELAKEGITKYDLGREKFL
Thermincola potens                        GHAIDNTLQDILTRWRRMQGYNALWLPGTDHAGIATQAKVEEQLAKEGLSKYDLGRDKFL
Thermofilum pendens                       GNALNFSYIDFVARYKRMRGYNVLFPQGWDCHGLPTEVRVEKA---VGKRKSEMDPNEFL
Thermoprotei archaeon bit383              GNALNFSYIDFVARYKRMRGFNVLFPQGWDCHGLPTEVRVEKV---TGKKKSEFDPQEFI
Crenarchaeota archaeon bit341             GNALNWCYIDFVARYKRMKGFNVHFPQGWDCHGLPTEVRVEQT---FKIRKNDLPPEQFV
C.Verstraetearchaeota archaeon bit357     GNALNWTYIDAVARYMRMKGYNVHFPQGWDCHGLPTEVRAERT---MNVNKREVDYEKFR
C.Bathyarchaeota archaeon bit301          GNVLNWTYFDIVARFKRMCGYNVLFPQGWDCHGLPTEVETEEK---YGIKKTDVPPAEFR
Hadesarchaea archaeon bit408              GTALNWTYFDIVARYKRMRGFNVLFPQGWDCHGLPTEVQVEKR---YGIRKGDLPPEKFR
Methanothrix thermoacetophila             GNALNWCYIDFVARYKRMRGYNVMFPQGWDCHGLPTEVKVEET---YHITKNQVPREEFR
Methanoplanus limicola                    GNAFNWCYIDFIARYKRMNGYNVMFPQGWDCHGLPTEVKVEET---HGITKNDVPRTEFR
Geoglobus acetivorans                     GNTLNWCYIDFIARYRRMKGYEVMFPQGWDCHGLPTEVKVEEK---YGIKKNDVPRERFR
Ferroglobus placidus                      GNALNWCYIDFIARYKRMKGYEVMFPQGWDCHGLPTEVKVEEL---YGIKKNDVPREKFR
Archaeoglobus fulgidus                    GHALNWCIIDFIARYKRMNGYEVMFPQGWDCHGLPTEVKVEEK---YGIKKGDIPRDEFR
Halococcus saccharolyticus                GNALGWCYMDFAARYHRLQGENVSFPQGWDCHGLPTEVKVEEN---QDIRRTEVSREEFR
Halalkalicoccus jeotgali                  GNALGWCYMDFAARFQRLQGKDVLYPQGWDCHGLPTEVKVEEN---QGIHRTDVSREEFR
Methanopyrus kandleri                     GHVLNWTYMDVVARYKRMCGYDVFFPQGWDCHGLPTEVKVEEI---HGITKRDAPRREFR
                                          *  .     :   *: *: * :  :    *  .:  :   :                .*
```

```
                                                          130       140       150       160       170       180
                                                 .........|.........|.........|.........|.........|.........|
Thermococcus sp. 2319x1                          KKCIEWTWQAIEAMRNQFIRIGYSADWDL--------------EYHTMDDDYKALVQKSL
Thermococcus litoralis                           KKCIEWTWQAIEAMRNQFIRIGYSADWDL--------------EYHTMDDDYKALVQKSL
Thermococcales archaeon bit391                   KKCIEWTWQAIESMRNQFIRIGYSADWDL--------------EYHTMDDDYKALVQKSL
Thermococcus sibiricus                           KKCIEWTWQAIEAMRNQFIRIGYSADWEL--------------EYHTMDDEYKALVQKSL
Thermococci archaeon B45 G15                     KKCIEWTWQAIEAMRTQFIRIGYSADWEL--------------EYHTMDDDYKALVQKSL
Palaeococcus pacificus                           KKCIEWTWEAIEAMRNQFIQIGYSADWDL--------------EYHTMDEEYKALVQKSL
Thermococci archaeonB89 G9                       KKCIEWTWEAIEAMRNQFIRIGYSADWEL--------------EYHTMDDEYKALVQKSL
Thermococcus kodakarensis                        QKCIEWTWQAIEAMRNQFIRIGYSADWDL--------------EYHTMDDWYKAAVQKSL
Pyrococcus furiosus                              KKCIEWTWQAIEKMRAQFIRIGYSADWDL--------------EYHTMDDWYKAAVQKSL
Thermococci archaeon B61 G1                      EACKKWTEYSIERMSRQLKSVGYFPDWNN--------------VYVTMDRDYWRRVQYSL
Thermococci archaeon B48 G16                     NLCLEETKLGAKKYRNIWTKLGISVDWDL--------------SYSTINKHCQRLAQISF
Thermococci archaeon B54 G1                      NLCLEETKLGAKKYRNIWTKLGISVDWDL--------------SYSTINKHCQRLAQISF
Euryarchaeota archaeon bit403                    KLCSEFANEKIDEMKTQFILLGESMDPSV--------------YYQTDEENYRRLTQLSF
Mahella australiensis                            ERAWQWREKYGRRITAQLKRLGVSCDWSR--------------ERFTMDEGCSRAVIEVF
Thermincola potens                               ERVWQWKEFYHNRIATQLRSLGSSCDWER--------------ERFTMDEGCSRAVQKVF
Thermofilum pendens                              RLCREYTLKWIESMKAALKGLGLSIDWST--------------EYKTMDPDYWRRTQLSF
Thermoprotei archaeon bit383                     RLSKEFTLKWIDAMKKALKALGLSIDWST--------------EYRTMDPDYWRRTQLSF
Crenarchaeota archaeon bit341                    EMCRKLTGEYIAKMKQSMNLLGISSDWAL--------------EYKTMDPSYYKLTQLSF
C.Verstraetearchaeota archaeon bit357            RICFELTEKWIEQMKKTMRRMGYSIDWSL--------------EYRTMDPDYWRRTQLSF
C.Bathyarchaeota archaeon bit301                 KLCESFVRKYIDLMKKAIIRLGCSIDWRL--------------EYQTMDPDYWKKTQLSF
Hadesarchaea archaeon bit408                     ELCIKLTEENITHMREEMNSLGFSIDWST--------------EYRTMDPDYYRRTQLSF
Methanothrix thermoacetophila                    RLCEEMTAQAIERMRRTITRLGISTDWSN--------------EYITMKPEYYVKTQRSF
Methanoplanus limicola                           EMCRELTLENIEKMRKSMRRCGFSNDWSN--------------EYITMLPEYYGKTQLSF
Geoglobus acetivorans                            ELCVEFTEGNIEKMRKTMRRVGFSIDWSK--------------EYITMYPEYYRKTQVSF
Ferroglobus placidus                             ELCVEFTERNIKRMKETMKKLGLSIDWSK--------------EYVTMYPEYYRKTQVSF
Archaeoglobus fulgidus                           RLCVEFTEENIAKMRETARRMGYSIDWSK--------------EYITMYPEYYSKTQLSF
Halococcus saccharolyticus                       ELCIEHTEGQIDAMKETMGRLGFSQDWSQ--------------EYRTMDSEYWGKTQRSF
Halalkalicoccus jeotgali                         ELCVEHTESQIDAMKETMGLLGFSQDWDH--------------EFRTMDPSYWGKTQRSF
Methanopyrus kandleri                            KLCEELTLENIRKMREQLIQLGCSIDWWTDCIDYENEELKELGSYVTMDPDYIRRSQYGF
                                                 .   .                *   *                    *            :
```

```
                                            190       200       210       220       230       240
                                    .........|.........|.........|.........|.........|.........|
Thermococcus sp. 2319x1             LEFYKKGLLYQDKHPVYWCPRCRTSLAKAEVGYVEEDGYLYYIKLPIAGEDDYIPIATTR
Thermococcus litoralis              LEFYKKGLLYQDKHPVYWCPRCRTSLAKAEVGYVEEDGYLYYIKLPIAGENDYIPIATTR
Thermococcales archaeon bit391      LEFYKKGLLYQDKHPVYWCPRCRTSLAKAEVGYVEEDGYLYYIKLPIAGEDDYIPIATTR
Thermococcus sibiricus              LEFYKKGLLYQDKHPVYWCPRCKTSLAKAEVGYVEEDGYLYYIKLPIAGEEDYIPIATTR
Thermococci archaeon B45 G15        LEFYKKGLLYQDKHPVYWCPRCKTSLAKAEVGYVEEDGYLYYIKLPIVGEEDYIPIATTR
Palaeococcus pacificus              LEFYKKGMLYQDKHPVYWCPRCRTSLAKAEVGYVEEDGYLYYIKLPIAGEDDYIPIATTR
Thermococci archaeonB89 G9          LEFYKKGMLYQDKHPVYWCPRCRTSLAKAEVGYVEEDGYLYYIKLPIAGEDDYIPIATTR
Thermococcus kodakarensis           IEFYKKGMLYQAEHPVYWCPRCRTSLAKAEVGYVEEDGFLYYIKLPLADGSGHVPIATTR
Pyrococcus furiosus                 LDFYKKGLIYREEHPVYWCPRCRTSLAKAEVGYVEEEGYLYYIKLPLADGSGYIPIATTR
Thermococci archaeon B61 G1         LKFHEKGLIYIKEHPVHFCPSCESAIAKAEINYREEEGYLNYIKFKLEDGE-TIEIATTR
Thermococci archaeon B48 G16        IELYKMGRLERKKEPVIWCPHCRTAIAQAELEDKEEDTFLNTIIFKTDDGK-DLPIATTR
Thermococci archaeon B54 G1         IELYKMGRLERKKEPVIWCPHCRTAIAQAELEDKEEDTFLNTIIFKTDDGK-DLPIATTR
Euryarchaeota archaeon bit403       IELYTKGLIYKGEHPINWCPRCMTALADAEIEYKNRETKLNYIKFKLEDED--VIIATTR
Mahella australiensis               VRLYEKGLIYRGDRIINWCPECKTALSDAEVEYEEEHGHLWHIRYPFTDGSGYMVVATTR
Thermincola potens                  VDLYKKGLIYRDNYIINWCTNCRTTISDIEVEHTDQAGHFWHIRYPVKDSDEYVYLATTR
Thermofilum pendens                 VLMYNKGLIYRGEHPVIWCPRCETAIAEAEVEYEERDRPLYYFKFGVEGTGEELVVASTR
Thermoprotei archaeon bit383        VLMYKKGLAYRAEHPVIWCPRCETAIAEAEVEYVEKDRPLYYFKFKVKETGEDLIIASTR
Crenarchaeota archaeon bit341       IELFKSGHLYRGEHPVNWCPRDETAIAEAEVVYQERKGTLHYMRFGHA--NEHIEIASTR
C.Verstraetearchaeota archaeon bit357 IILYRKGLVYRAEHPVLWCPRCETAIAEAEVEYVERKTKLYYVKFRRSGSGE-VIVATTR
C.Bathyarchaeota archaeon bit301    IILYDKGYIYRGTHPVNWCPRCSTAIADAEVEYESREGRLHYIRFRMKGGGS-IPIATTR
Hadesarchaea archaeon bit408        VQLYKRGLIYRGEHPVNWCPRCETAIADAEVEYEDREAKLSYIKFKLHGEDEYLTIATTR
Methanothrix thermoacetophila       VQMYEKGMIYREDHPVNWCPRCATAIAFAEVEYDTRTTTLNYMRFESD-HG-CLEIATTR
Methanoplanus limicola              LRMFENGDVYQSEHPVNFCTRCETAIAFAEVSYEDRTTKLNFFDFDG------VEIATTR
Geoglobus acetivorans               VRMYRKGLIYRGEHPVIHCPRCETTIALAEIEYKSGKTKLNYIKFDDD-----VIIATTR
Ferroglobus placidus                VRLYEKGWIYKGYHPVIVCPRCETTIALAEIEYKEGKTKLNYIKFDEN-----VIIATTR
Archaeoglobus fulgidus              VRMYNKGLIYRDYHPVVFCPRCETTIALAEIEYRQGKTKLNYIKFDDD-----VIIATTR
Halococcus saccharolyticus          VEMAEKGYVHRDEHPVNWCPRCETAIADAEVENIDTEGTLSTVRFSGVDND-DIEIATTR
Halalkalicoccus jeotgali            VEMHSSEYVYRDEHPVNWCPRCETAIADAEVENEDREGTLFSIRFAGVENA-PIEIASTR
Methanopyrus kandleri               LELLEKGYAYREEHPVNWCPRCETAIAFAEVEYVTRETYLNYIEFPVADGDGSVTIATTR
                                    : :            :  *    ::::  *:        :  .          : :*:**
```

```
                                                         250       260       270       280       290       300
                                                .........|.........|.........|.........|.........|.........|
Thermococcus sp. 2319x1                         PELMPACVAVFVHPEDERYKGKVGKKVKLPIFER---------EVPILADEDVDPNFGTGA
Thermococcus litoralis                          PELMPACVAVFVHPEDERYKDKVGKKVKLPIFER---------EVPILADEDVDPNFGTGA
Thermococcales archaeon bit391                  PELMPACVAVFVHPEDERYKDKVGKKVKLPIFER---------EVPILADEDVDPNFGTGA
Thermococcus sibiricus                          PELMPACVAVFVHPEDERYKNKVGKKVKLPIFER---------EVPIIADEDVDPEFGTGA
Thermococci archaeon B45 G15                    PELMPACVAIFVHPEDERYKNKVGKKVKLPIFER---------EVPIIADEDVDPEFGTGA
Palaeococcus pacificus                          PELMPACVAVFVHPDDERYKDKVGKKVNLPIFER---------EVPIIADKDVDPSFGTGA
Thermococci archaeonB89 G9                      PELMPACVAVFVHPDDERYKDKVGKKVRLPIFER---------EVPIIADEDVDPEFGTGA
Thermococcus kodakarensis                       PELMPACVAVFVHPEDERYKHVVGKKVKLPIFER---------EVPVLADEDVDPSFGTGA
Pyrococcus furiosus                             PELMPACVAVFVHPDDERYKHLVGKKVKLPIYER---------EVPILADEDVDPNFGTGA
Thermococci archaeon B61 G1                     PELLPACVAVAVNPEDDRYKDLPGRKAIVPIFNQ---------KVEIFSDRAVDPEFGTGI
Thermococci archaeon B48 G16                    PEFLASCVSVVVHPFDERYKYLIGKYAIVPIFGN---------RVKIISDNRVDKDFGTGI
Thermococci archaeon B54 G1                     PEFLASCVSVVVHPFDERYKYLIGKYAIVPIFGN---------RVKIISDNRVDKDFGTGI
Euryarchaeota archaeon bit403                   PELLCTCQLVAIHPDDFRAEKLKGKKLKTPLFEK---------EVRIVGDKNVDPNFGSGI
Mahella australiensis                           PETMLGDTAVAVNPNDERYKDVIGKTVILPLINR---------EISVIADEYVDMEFGTGA
Thermincola potens                              PETMLGDTAVAVHPADERYKHLVGRSVILPLVGR---------EIPVIADEYVDPEFGTGV
Thermofilum pendens                             PELLASCVAVAVNPSDERYKHLVGKNAVVPIYGR---------KVPIIADEAVDKDFGTGA
Thermoprotei archaeon bit383                    PELLASCVAVAVHPEDERYRNIVGKKAIVPIYER---------EVPIITDPEVDKEFGTGA
Crenarchaeota archaeon bit341                   PELLSACVGIAVHPKDEKYKLVVGKTIAVPIFGQ---------QARVISDDEVDPRYGTGA
C.Verstraetearchaeota archaeon bit357           PELIPACVGVLVNPDDERYRDLIGKVLITPIFER---------EVKVYADREVDPTFGTGV
C.Bathyarchaeota archaeon bit301                PELIPACVAVAVHPDDERYSEYVGGTVEVPLVGR---------EVPIIEDEAVDPDFGTGA
Hadesarchaea archaeon bit408                    PEYLPACVVVAVYPGDERYGKYVGKKVEVPPFGQ---------VVELIEDREVDPEFGTGV
Methanothrix thermoacetophila                   PELLPACVAVAVNPNDERHIGFVGKSVKVPLFDY---------EVPVLSDPAVDPSFGTGV
Methanoplanus limicola                          PELLAACVAVAVHPDDNRYSEISGKTLKVPIFGH---------GVKVIEDEAVDPSFGSGA
Geoglobus acetivorans                           PELIPACVAIAVHPEDERYRDVVGKTVRVPGTGH---------EVKVIADEDVDPEYGTGM
Ferroglobus placidus                            PELIPACVAVAVHPEDERYKDLVGKTVVVPISGK---------EVKIIADEEVDPNYGTGM
Archaeoglobus fulgidus                          PELIPACVAIAVHPDDERNKHLIGKKVRVPTTPY---------EVEVIADEEVDPEFGTGV
Halococcus saccharolyticus                      PELLAACVGMAVDPDDERYADRVGDTFEVPLFGQ---------EVELIADEEVDGDFGTGA
Halalkalicoccus jeotgali                        PELLGACVAIAVSPDDERYEDRVGESFEVPLFGQ---------EVELISDEEVDSEFGTGA
Methanopyrus kandleri                           PELLPACVAVAVHPDDDRYSDLVGKKLVVPLHERFGDRDTPWEVPVIADEEVDPEFGTGI
                                                ** :     : : * * :     *     *               :  *  **  :*:*
```

```
                                                310       320       330       340       350       360
                                        .........|.........|.........|.........|.........|.........|
Thermococcus sp. 2319x1                 VYNCTYGDEQDVVWQKRYNLPVIIAIDENGRMNENAGKYKGLTAEEAREAIARDLEEMGL
Thermococcus litoralis                  VYNCTYGDEQDVVWQKRYNLPVIIAIDENGRMNENAGKYKGLTAEEAREAIAKDLEEMGL
Thermococcales archaeon bit391          VYNCTYGDEQDVVWQKRYNLPVIIAISEDGRMNENAGKYKGLTTEEARKAIAKDLEEMGL
Thermococcus sibiricus                  VYNCTYGDEQDVLWQKRYNLPVIITINEDGTMNENAGKYKGLTAEQAKEAIVKDLEEMGL
Thermococci archaeon B45 G15            VYNCTYGDEQDVVWQKRYNLPVIITINEDGTMNENAGKYKGLTAEQAKEAIVKDLEKMGL
Palaeococcus pacificus                  VYNCTYGDEQDVVWQKRYKLPVIITINEDGTLNENAGKYKGLTSEEARDAIAKDLEELGL
Thermococci archaeonB89 G9              VYNCTYGDEQDVVWQKRYNLPVIIAIDEDGTLNENAGKYKGLKAEDAREKIAEDLEKLGF
Thermococcus kodakarensis               VYNCTYGDEQDVVWQKRYNLPVIIAINEDGTMNENAGPYAGLKTEEARKKIAEDLEKMGL
Pyrococcus furiosus                     VYNCTYGDEQDIVWQKRYNLPVIIAINEDGTMNENAGPYAGLKIEEARKKIAEDLEKMGL
Thermococci archaeon B61 G1             VMICTFGDEQDVKWAYERGLPVVKAINQRGKLTKVAGKYKGMGILEAREAIISDLEKQGY
Thermococci archaeon B48 G16            VMICTFGDKTDIEWWKDYNLDLVISINKDGTLNEKAGKFKGMELKEARKAVLKELESKNL
Thermococci archaeon B54 G1             VMICTFGDKTDIEWWKDYNLDLVISINKDGTLNEKAGKFKGMELKEARKAVLKELESKNL
Euryarchaeota archaeon bit403           VMICSIGDKDDLEWIFKYKLKIEKGIDEEGRMTGLCGKYRGMKVSDARKAIIDDMKKQGA
Mahella australiensis                   VKITPAHDPNDFEVGIRHNLPVIRVMADDGHMNENAGRYNGMERYEARRAVLADLEAQGF
Thermincola potens                      VKITPAHDPNDFEVGKRHNLPEITVMNKDGTMNEEAGIYRGLDRYECRKRIVEDLEKGGF
Thermofilum pendens                     VMVCTYGDKTDVKWQKKYNLPVIISITEQGTMNDNAGPLKGLKVEDARKKIVEMLKENGL
Thermoprotei archaeon bit383            VMICTYGDKTDVKWQKRYNLPIIISITDKGFMNENAGPLKGLRIEEARKKMVELLRENGL
Crenarchaeota archaeon bit341           VMICTFGDKTDVRWQAKYHLPVIKALTENGRLSVDDPRFKGLKAEEARRKIVAELQASGL
C.Verstraetearchaeota archaeon bit357   VMICTFGDKTDVRWQKKFNLPIIKAIDERGVMTDAAGEFKGLTIENCREKIVLKLREEGY
C.Bathyarchaeota archaeon bit301        VMICTYGDKDDVKIVAKHGLPVIMILDEEGRLNEKAGKYKGLTVQEARKAIVEDLRSEGY
Hadesarchaea archaeon bit408            VMVCTFGDKTDVRWVKRHKLPVVKLIDEKGNMSEAAGKYTGMTLEECKSKIVDDLRKAGL
Methanothrix thermoacetophila           VMICTFGDKQDVRWWVEHKLPLRQAIDREGRLTEIAGKFGGMSITEAKKAIVDEMLSRGI
Methanoplanus limicola                  VMICTFGDKQDVHWWKKHNLDLRKAIDLKGRMTAIAGPYAGMTSQECRNGILSEMAEKGI
Geoglobus acetivorans                   VMICTFGDRQDVRWWKKHRLELRDVLTRDGRLNEKAGKYAGLNVREAREKILEDLEKEGR
Ferroglobus placidus                    VMICTFGDKQDVRWWKKHGLELRQVLTRDGKLNEKAGKYAGLTVSEAREKIIEDFKKEGR
Archaeoglobus fulgidus                  VMICTFGDRQDVKWWKKHKLELRNIVGRDGRLNEKAGRYAGMTIPEAREAILEDLKKEGK
Halococcus saccharolyticus              VMICTFGDKQDVDWWAEYDLDLRPVVTEDGRLAEDVPEFGGLAIDEAKAKISTALQKEGY
Halalkalicoccus jeotgali                VMICTFGDKQDVDWWAEYDLPLRTVLTEDGRLNERAGEFEGLEIDEAKTEIAAALDEAGN
Methanopyrus kandleri                   VMICTFGDKQDVAWVKRHDLPIVRAIDEQGKMTEVAGEFAGMEVEEARAAIVEALKEEGY
                                        *      *  *.       *     :   * :        *:   :.:  :   :   .
```

```
                                                        370       380       390       400       410       420
                                                .........|.........|.........|.........|.........|.........|
Thermococcus sp. 2319x1                         LYKKEKVHHRVLRHTERSSCMAPIELLPKKQWFIKVKDFTDEIVKVAE--QIKWYPEDMF
Thermococcus litoralis                          LYKKEKVHHRVLRHTERSSCMAPIELLPKKQWFIKVKDFTDEIVKVAE--QIKWYPEDMF
Thermococcales archaeon bit391                  LYKKEKVHHRVLRHTERSSCMAPIELLPKKQWFIKVKDFTDEIVKVAE--QIKWYPEDMF
Thermococcus sibiricus                          LYKKKQVHHRVLRHTERSSCRAPIELLPKKQWFIKVKELTDEIVKVSE--QIRWYPPDMF
Thermococci archaeon B45 G15                    LYKKKQVHHRVLRHTERSSCRAPIELLPKKQWFIKVKELTDEIVKVSE--QIRWYPPDMF
Palaeococcus pacificus                          LYQKKKVHHRVLRHTERSSCKAPIELLPKKQWFIKVKDFTEDIVKVAE--QINWYPEDMF
Thermococci archaeonB89 G9                      LYQKKKVHHRVLRHTERSSCKASIELLPKKQWFIKVKDFTDEIIKVAE--QINWYPEDMF
Thermococcus kodakarensis                       LYKKEKIRHRVLRHTERSSCMAPIELLPKKQWFIKVKDFTDEIVKVAE--QINWYPPDMF
Pyrococcus furiosus                             LYKKEKITHRVLRHTERSSCMAPIELLPKKQWFIKVREFTDDIVEVAK--KINWYPEDMF
Thermococci archaeon B61 G1                     LVKREKIVHQVA---VHDKCGTPIEFVPTLQWFIKVRQFKEDIIGAAM--SMKWIPDYML
Thermococci archaeon B48 G16                    LIKKEKLRHAVN---THERCGTSVEYLVEDQWFVKILDLKDVWIKQAD--KMNWYPDFMK
Thermococci archaeon B54 G1                     LIKKEKLRHAVN---THERCGTSVEYLVEDQWFVKILDLKDVWIKQAD--KMSWYPDFMK
Euryarchaeota archaeon bit403                   LIKQEKLEQSIG---TCWRCHTPVEFLVMPQWFLKTVQFKEKMLKEIE--KLDWKPRFMK
Mahella australiensis                           IDKIENHVHNVG---HCYRCSTIVEPIVSKQWFVKMEPLAEPAIEAVRDGRIQFIPQRFA
Thermincola potens                              LVKVEDHTHSVG---HCYRCGTVIEPLVSKQWFVKMKPLAEPAIQAAKEERVRFIPERFT
Thermofilum pendens                             LVKVESIRSTVG---TCWRCHTPVEIIPKKQWFVRSTALNEKVLEEGR--KVNWVPSYMY
Thermoprotei archaeon bit383                    LVKVETVRSSVG---TCWRCHTPVEILPKKQWFIKSRELAEKVLEEAR--KVQWVPPYMY
Crenarchaeota archaeon bit341                   LTKTEPITQNIG---TCWRCQTPVEIIARPQWFMKTRDLTQKVVEWAG--KLDWIPSFSK
C.Verstraetearchaeota archaeon bit357           IVREEHIKQNVG---TCWRCHTPVEILAKEQWFIKTRELAEDVVNWAG--RIKWIPKWAE
C.Bathyarchaeota archaeon bit301                LEKVESLRQEVG---VCWRCKTPVEILERTQWFMKTRVLTERVEQEAY--KIAWYPDHMR
Hadesarchaea archaeon bit408                    LVKQEKLAQSVG---ACWRCNTPVEILTKPQWFMRVLDLKDKVIEGAK--RVRWVPEHMR
Methanothrix thermoacetophila                   IYRQEPLEQNVG---LCWRCKTPIEILSERQWFVRIY--PDVIIKTAD--EIEWVPEHMK
Methanoplanus limicola                          LKRQEDLEQRVG---ACWRCKTPIEILSERQWFVKVH--NDEILKAAD--KIRWTPEHMK
Geoglobus acetivorans                           LIKQEEVDHNVG---VCWRCKTPVEIVPEKQWFVKIE--KEKILEAAR--RVKWVPEHML
Ferroglobus placidus                            LIKVEEVDHKVG---VCWRCKTPVEIIPEEQWFVKVD--KERVLEAAR--KIKWIPEHMY
Archaeoglobus fulgidus                          LLKQVEIDHNVG---TCWRCKTPVEIIPAEQWFVKVE--KEKILEAAK--RIKWVPEHMY
Halococcus saccharolyticus                      LQDEEPIEQSVG---ACWRCDTPIEILSKEQWFVRVD--REEILGKAQ--EIDWIPEHMY
Halalkalicoccus jeotgali                        LQGEEPVEQSVG---CCWRCDTPIEILSKEQWFIRVD--GEEILENAR--EIEWIPEHMY
Methanopyrus kandleri                           LVKREKITQNVG---VCWRCKTPIEILVKEQWFVKVRELAEDVKEAAR--KMVWIPEHMR
                                                :         :        * : :* :   ***::     :          : : *
```

```
                                                  430       440       450       460       470       480
                                         .........|.........|.........|.........|.........|.........|
Thermococcus sp. 2319x1                  LRLKDWAESMDWDWVISRQRVFGTPIPFWVCKDCGEIIPAREEDLPVDPRFDKPPVEKCP
Thermococcus litoralis                   LRLKDWAESMDWDWVISRQRVFGTPIPFWVCKDCGEIIPAREEDLPVDPRFNKPPVEKCP
Thermococcales archaeon bit391           LRLKDWAESMDWDWVISRQRVFGTPIPFWVCRDCGEIIPAREEDLPVDPRFDKPPVDKCP
Thermococcus sibiricus                   LRLKDWAESMDWDWVISRQRVFGTPIPFWVCKDCGEIIPAREEDLPVDPRFNGAPVDKCP
Thermococci archaeon B45 G15             LRLKDWAESMDWDWVISRQRVFGTPIPFWVCKDCGEIIPAREEDLPVDPRFNGPPINKCP
Palaeococcus pacificus                   LRLKDWAESMDWDWVISRQRVFGTPIPFWVCKECGHIVPAKEEDLPVDPRFDKPPVERCP
Thermococci archaeonB89 G9               LRLKDWAESMDWDWVISRQRVFGTPIPFWVCRDCGEIIPAREEDLPVDPRFDKPPVERCP
Thermococcus kodakarensis                LRLKDWAESMDWDWVISRQRVFGTPIPFWVCDN-GEIILPNEEDLPVDPRFEKPPRKC--
Pyrococcus furiosus                      LRLKDWAESMDWDWVISRQRVFGTPIPFWICKN-GHIIPAREEDLPVDPRFDKPPVEKCP
Thermococci archaeon B61 G1              QRLIDWAESLDWDWVISRQRIWGTPIPFWYCEKCHYIIAPREEELPLDPTEREPPVKECP
Thermococci archaeon B48 G16             KRYISWVENLRWDWCISRQRYYGVPFPLWYCKKCGNVVLPEKEDLPVDPLKDKPKK-PC-
Thermococci archaeon B54 G1              KRYISWVENLRWDWCISRQRYYGVPFPLWYCKKCGNVVLPEKEDLPVDPLKDKPKK-PC-
Euryarchaeota archaeon bit403            RRLQDWINSLSWDWVISRQRYFATPIPIWECKKCGNIVVAKKNECYVDPTIMD---KECP
Mahella australiensis                    NIYFNWMENIK-DWCISRQLWWGHRIPAYYCHVCGNIMVLS-----------QRP-HTCD
Thermincola potens                       KIYLNWMENIR-DWCISRQLWWGHRIPVWYCQDCGEVICEL-----------DPP-EKCP
Thermofilum pendens                      KRLENWVLSLDWDWVISRQRLFATPIPVYYCKDCGAELVVPPEKLPIDPRFDPPPFEKCP
Thermoprotei archaeon bit383             KRLENWVESLDWDWVISRQRLFATPFPVYYCKKCGHEIVVEPEHLPIDPRKDPPPVKVCP
Crenarchaeota archaeon bit341            QRLIDWAESLDWDWVVSRQRIFATPIPVWYCTKCESTIIPNEESLPVDPRKDKSPGEECP
C.Verstraetearchaeota archaeon bit357    KRLIDWVKALDWDWVISRQRIFATPIPVWYCKKCGKIIVAEEEWLPVDPRRDRPRINRCP
C.Bathyarchaeota archaeon bit301         NRLIDWAKSLNWDWVISRQRVFATPIPIWYCAKCGEVILAKPEWLPIDPKLEKPKIDKCP
Hadesarchaea archaeon bit408             QRLIDWAESMDWDWVISRQRIFATPIPAWYCTGCREPIVAEEKQLPVIPAKDKPLVKRCP
Methanothrix thermoacetophila            LRLKNWTGTMEWDWCISRQRVFATPIPAWYCKRCGEVMVAKEEWLPLDPTKTQPPV-SC-
Methanoplanus limicola                   MRLENWASQMEWDWCISRQRIFATPIPVWFCNKCGEVMLPDEEDLPVDPTVDRPKR-PCP
Geoglobus acetivorans                    SRLESWVESMDWDWVISRQRVFATPIPVWYCRNCGEVVVAKEEWLPVDPTRDKPRE-PCP
Ferroglobus placidus                     LRLEDWVESMEWDWVISRQRVFATPIPVWYCENCGETIVAKEEWLPVDPTKDQPKE-PCP
Archaeoglobus fulgidus                   SRLESWVQSMEWDWVISRQRIFATPIPAWYCKNCGEVVVAKEEWLPVDPTATQPPE-PCP
Halococcus saccharolyticus               GRLEEWTEGMEWDWVISRQRVFATPIPAWSCDECDHWHIATLDELPAEPTEDDPDI-ECP
Halalkalicoccus jeotgali                 ARLEEWTEGMEWDWVISRQRVFATPIPAWFCEECEYVHVADTEALPVKPTSDGPDI-ACP
Methanopyrus kandleri                    KRLEDWTESMDWDWCISRQRIFATPIPVWYCKECGEVIPAEKDQLPVDPTRDDPPVDECP
                                             .*   :  ** :***  :.  :* : *
```

```
                                                490       500       510       520       530       540
                                        .........|.........|.........|.........|.........|.........|
Thermococcus sp. 2319x1                 KCGSTNIEGVKDVLDCWVDSSITPLVISKWQK-------NERWFKHNFPTSLRPQGTDII
Thermococcus litoralis                  KCGSTNLEGVKDVLDCWVDSSITPLVISKWQR-------DEKWFKHNFPTSLRPQGTDII
Thermococcales archaeon bit391          KCGSTNLEGVKDVLDCWVDSSITPLVISKWQR-------DEKWFKHNFPTSLRPQGTDII
Thermococcus sibiricus                  KCGSTNLEGVKDVLDCWVDSSITPLVISKWQK-------DEKWFSHNFPTSLRPQGTDII
Thermococci archaeon B45 G15            KCGSTNLEGVKDVLDCWVDSSISPLVVSKWQK-------DEKWFSHNFPTSLRPQGTDII
Palaeococcus pacificus                  KCGG-ELEGAKDVLDCWVDSSISPLAITKWKR-------DEEWFKINFPTALRPQGHEII
Thermococci archaeonB89 G9              KCGSTNLEGAKDVLDCWVDSSVSPLAITKWGK-------DERWFKINFPTALRPQGHEII
Thermococcus kodakarensis               SDGS-EPKPVTDVLDCWVDSSITPLIITKWHEAIKGDEEGKKWFEHNFPTALRPQGTDII
Pyrococcus furiosus                     VCGA-EIEPVTDVLDCWVDSSITPLIITKWHEAIKGDEEAKKWFEHNFPTALRPQGTDII
Thermococci archaeon B61 G1             SCGSRRISPARDVCDCWVDSSITPLVITGWPN-------EREKFEKLYPVSIRPQGYEII
Thermococci archaeon B48 G16            KCGSTDFIPEEDVMDTWFTSSLTPQIIARWEC-------EDSLMDKIYPNSLRPQAHDII
Thermococci archaeon B54 G1             KCGSTDFIPEEDVMDTWFTSSLTPQIIARWEC-------EDSLMDKIYPNSLRPQAHDII
Euryarchaeota archaeon bit403           ECRT-TMKGCEDVFDTWMDSSISPLFNTCWER-------NPDLFKKLYPMSLRPQAHDII
Mahella australiensis                   KCGSDDIEQDPDVLDTWFSSALWPFSTLGWPD-------RTLELDYFYPTDVLVTAYEII
Thermincola potens                      KCGSARLEQDPDVLDTWFSSGLWPFSTLGWPE-------RKPELEYFYPTSVLVTGRDII
Thermofilum pendens                     KCGSKNIVPERDVMDTWMDSSITAAVHAGWPD-------NFD--ERLFPADLQPNGYDII
Thermoprotei archaeon bit383            KCGSTELVGETDVMDTWMDSSITAAVHAGWPD-------NMD--ERLFPADLQPNGYDII
Crenarchaeota archaeon bit341           RCGNRELRGETDVLDTWMDSSITAAVHAGWPN-------KSELFNRLFPADLQPNGLDIV
C.Verstraetearchaeota archaeon bit357   RCGGEDFEGERDVMDTWMDSSITCAVHAGWPD-------KPEVFKRLFPADLQPNGYDII
C.Bathyarchaeota archaeon bit301        KCGSTEFKPETSVLDTWFDSSITCAVHAGWPD-------KE-DWRRFFPADVHPSGADII
Hadesarchaea archaeon bit408            KCGGESFEPERDVLDTWMDSSITIAVHAGWPE-------LD---RRLFPADLQPNGTDII
Methanothrix thermoacetophila           SCGSNEFEPEEDVLDTWMDSSISALHVTGWLS-------RE--DPR-YPAQLRPQGHDII
Methanoplanus limicola                  KCGCTDFSGETDVLDTWMDSSISVLNITGWDG-------KS--VPEYFPAQIRPQGHDII
Geoglobus acetivorans                   KCGGNEFDGEDDVLDTWMDSSITSLAITGWPD-------D---QKE-YPTHLRPQGHDII
Ferroglobus placidus                    KCGSTKFRGEDDVLDTWMDSSITPLAIVGWPE-------E---LKE-YPTHLRPQGHDII
Archaeoglobus fulgidus                  KCGSTEFRGETDVLDTWMDSSITPLMICGWPS-----------LKE-YPTHLRPQGHDII
Halococcus saccharolyticus              DCGADAWTGETDVMDTWMDSSISALHAAGWPD-------A-----EFTPVQLREQGHEII
Halalkalicoccus jeotgali                ECGSEDWRGETDVMDTWMDSSISALHVAGWPD-------E-----EFSPVQLREQGHDII
Methanopyrus kandleri                   KCGCSEFEPETDVMDTWMDSSITPLVITGWPD-------EEP----DLPVDLRPQGHDII
                                                   .* * *. *.:       *                  *  :   . :*:
```

```
                                                  550       560       570       580       590       600
                                         .........|.........|.........|.........|.........|.........|
Thermococcus sp. 2319x1                  RTWAFYTIFRTYMLT-GQKPWHDILINGMVAGPDGRKMSKSYGNVVSPEEVIPKYGADAL
Thermococcus litoralis                   RTWAFYTIFRTYMLT-GQKPWHDILINGMVAGPDGRKMSKSYGNVVSPEEVIPKYGADAL
Thermococcales archaeon bit391           RTWAFYTIFRTYMLT-GQKPWHDILINGMVAGPDGRKMSKSYGNVVSPEEVIPKYGADAL
Thermococcus sibiricus                   RTWGFYTIFRTYILT-GEKPWDDILINGMVAGPDGRKMSKSYGNVVSPEEVIPKYGADAL
Thermococci archaeon B45 G15             RTWGFYTIFRTYILT-GEKPWDDILINGMVAGPDGRKMSKSYGNVVSPEEVIPKYGADAL
Palaeococcus pacificus                   RTWAFYTIFRSYVLT-GQKPWHDIMINGMVAGPDGRKMSKSLGNVITPEEVIPKYGADAV
Thermococci archaeonB89 G9               RTWAFYTIFRSYVLT-GQKPWEDIMINGMVAGPDGRKMSKSLGNVITPEEVIPRYGADAV
Thermococcus kodakarensis                RTWAFYTIFRTWVLT-GEKPWHDILINGMVAGPDGRKMSKSYGNVVAPDEVIPKYGADAL
Pyrococcus furiosus                      RTWAFYTIFRTYMLT-GEKPWNDIVINGMVAGPDGRKMSKSYGNVVSPEEVIPKYGADAL
Thermococci archaeon B61 G1              RTWAFYTIFRCLMLT-GKPPFKEIVINGMVLGDDGRKMSKSYGNVVEPDEVVEKYGGDSL
Thermococci archaeon B48 G16             RTWLFYTVVKSYFHN-KSIPWYSIMLSGFGLDEEGKAMHKSKGNVIHPLEIVEKYSADAV
Thermococci archaeon B54 G1              RTWLFYTVVKSYFHN-KSIPWYSIMLSGFGLDEEGKAMHKSKGNVIHPLEIVEKYSADAV
Euryarchaeota archaeon bit403            RTWAFYTILRGILLT-NKLPFKTIMVDGFVLAPDGKPMHASLGNVIDPLKILEKYGSDAF
Mahella australiensis                    FFWVARMIFSSLEHM-GDIPFKYVLIHGIVRDDEGRKMSKSLGNGVDPLDVVDKYGADAL
Thermincola potens                       FFWVARMIFMAMEFM-KEVPFREVFIHGLVLDAQGRKMSKSLGNGVDPIEVIEKYGADTL
Thermofilum pendens                      RTWDYYLILRGVALF-GRSQFKTALINGMVRGTDGRMMHKSYGNYVAVQEVLEKYGADSF
Thermoprotei archaeon bit383             RTWDYYLILRGVALF-GKSQFKTALINGMVRGTDGRMMHKSYGNYVALLDVIEQYGADAF
Crenarchaeota archaeon bit341            RTWDYYLLVRSLALF-GRPPYKTLLINGMVKGTDGRMMHKSYGNYVVADEAIKKVGADAL
C.Verstraetearchaeota archaeon bit357    RTWDYYLIVRGLMLF-GMPQFKTALINGMVRGTDGRMMHKSYGNYVGADEAIEKYGADAL
C.Bathyarchaeota archaeon bit301         RTWAYYLMVRHLALF-DEKPFKACLINGMVLGTDGRKMSKSLGNYIATDEVFGKYGADAT
Hadesarchaea archaeon bit408             RTWDYYLLVRHLALL-GEVPYRTVLINGMVFGEDGRKMSKSLGNYIDTTMARGKYGTDAL
Methanothrix thermoacetophila            RTWAFYTILRSMALV-GVKPWETILINGMVLGEDGRKMSKSLNNFVIPEEVFEKNGADAL
Methanoplanus limicola                   RTWAFYTILRAGALTGGGHPWDEILVNGMVLGDDGFKMSKSRGNIIVPEEVLSKHGADAF
Geoglobus acetivorans                    RTWAFYTILRSIALV-GEIPWHEIVINGMVLGEDGRKMSKSLGNVISPEEVIEKYGADAL
Ferroglobus placidus                     RTWAFYTILRSIYLV-NQIPWYEIVINGMVLGEDGRKMSKSLGNYVEAEEVIEKYGADCL
Archaeoglobus fulgidus                   RTWAFYTILRSLALE-GQIPWYEIVINGMVFGEDGRKMSKSLGNVIVPEEVVEKYGVDAL
Halococcus saccharolyticus               RTWAFYTILRTAALE-GEKPWDEALINGMVLGTDGNKMSKSKDNSVAPEEVVKEHSADAF
Halalkalicoccus jeotgali                 RTWAFYTILRTAALE-EQIPWEQALINGMVFGEDGNKMSKSRGNFVQPEEVVEEHGADAF
Methanopyrus kandleri                    RTWLYYTTVRALVHA-DTEPFKEILINGMVFGEDGYKMSKSRGNVVEPTEVIEEYGADAL
                                           *     .           :   .: *:    :*  *  * .* :       . . *
```

```
                                                 610       620       630       640       650       660
                                        .........|.........|.........|.........|.........|.........|
Thermococcus sp. 2319x1                 RLWTAL-APPGEDHPFKWEIVDYNYRFLQKLWNIFRFAERHIKGFDYE-KYK----DSEL
Thermococcus litoralis                  RLWTAL-APPGEDHPFKWEIVDYNYRFLQKLWNIFRFAERHIKGFDYE-KYK----DLEL
Thermococcales archaeon bit391          RLWTAL-APPGEDHPFKWEIVDYNYRFLQKLWNIFRFAERHITDFDYE-KYK----DIEL
Thermococcus sibiricus                  RLWTAL-APPGEDHPFKWEIVDYNYRFLQKLWNIFRFAERHIKDFDYE-EYK----DIEL
Thermococci archaeon B45 G15            RLWTAL-APPGEDHPFKWETVDYNYRFLQKLWNIFRFAERHIEDFEYN-KYK----DIEL
Palaeococcus pacificus                  RIWTAL-APPGEDHPFKWETVDYNYRFLQKLWNVFRFAERHIKDFDYE-KHK----EIEL
Thermococci archaeonB89 G9              RIWTAL-APPGEDHPFKWETVDYNYRFLQKLWNIFRFAERHIKDFDYE-KHK----EIDL
Thermococcus kodakarensis               RLWTAL-APPGEDHPFKWETVDYNYRFLQKVWNIYRFAERHLENFDPA-S-A----PEEL
Pyrococcus furiosus                     RLWTAL-APPGEDHPFKWEIVDYNFRFLQKLWNIYRFAERHIKDFDYE-KYK----HIEL
Thermococci archaeon B61 G1             RQWVST-GALGKDVPFSWKEVKHGHKFLRKLWNIARFVKQNTSDYDGS-EYD----VSEL
Thermococci archaeon B48 G16            RWWSSS-VKLGDDLPYSEKDVVAGHRLCIKLWNASKLISSHLDEKIES---------SDL
Thermococci archaeon B54 G1             RWWSSS-VKLGDDLPYSEKDVVAGHRLCIKLWNASKLISSHLDEKIES---------SDL
Euryarchaeota archaeon bit403           RYYAAT-CAVGEDNSFREKDVIHGMRLCTKLWNVSSFVKKIIDGKKIVLE------KSRL
Mahella australiensis                   RFTLVTGNAPGNDMRFYWDKVEASRNFANKIWNAARFVLLNLNEQDIKPTS-----LDEL
Thermincola potens                      RFMLITGNTPGNDLRFHFERLDNTRNFANKIWNASRFVLMNLEDFKPGE-P-----PKDY
Thermofilum pendens                     RLWVALAAATGQDVRFSWDGVDYAHRFLVKVWNLARLASPFIEDVREVPL-------GNL
Thermoprotei archaeon bit383            RLWVATAASTGQDVRFSWEGLKYTKRYLTKIWNASRFAYIFIKDYKPKSK-------VEL
Crenarchaeota archaeon bit341           RQWAAAGGSTGYDIPFRWNELEHGKKFLTKLWNVTRFVLANTTEPLR--LKK----PSTL
C.Verstraetearchaeota archaeon bit357   RQWALTGGGTGSDIPFRWEDVDYGWRFLLKLWNASRFVSLHVKNYDGISG------AGSL
C.Bathyarchaeota archaeon bit301        RQWAAGGGATGSDIPFRWADVEYGRRFLRKLWNAARFISINLKDFDPEVE-------VDV
Hadesarchaea archaeon bit408            RQWAVIGASTGSDIPFSWKDVDFGYRFMRKFWNAARFAGPHLKAKISELDPK----KLKF
Methanothrix thermoacetophila           RQWAALGGSPGSDVMFQWKEIVAASRFQQKLWSIYRFAAPFASDT-----------DAPF
Methanoplanus limicola                  RQWSASGAATGQDIVFSWNDVIAAARFQTKMWNITRFVIMHLDKAEEGMIA-----DRPT
Geoglobus acetivorans                   RQWAAIGGIMGTDVIFSWKEVVAASRFQQKFWSILRFGMMHLDDYQPDESH-----RELL
Ferroglobus placidus                    RQWAAIGGVVGSDVAFSWKEIKSASRFQQKFWSLTRFSAMHLENYEPKDEH-----EQYL
Archaeoglobus fulgidus                  RQWAAS-GVIGDDIIFNWKDVIAASRFQQKFWSITRFTLMHISDYTPSEED-----KKLL
Halococcus saccharolyticus              RQAMALGGQPGSDIQFQPKEVTSASRFLTKLWNITRFASQHLDAESGGSAADDGAFDEAA
Halalkalicoccus jeotgali                RQAIALGGQPGSDIQFQSKEVTSASRFLTKLWNISRFSDNHLDEDTPDVD------APAY
Methanopyrus kandleri                   RYWAVSSGAPGSDVQYMTKTIKRGYRFAKKIWNVCRLAKDHIDDAPSV-EEV----EGDL
                                        *         * *  :    :    .   *.*.   :
```

```
                                                  670       680       690       700       710       720
                                         .........|.........|.........|.........|.........|.........|
Thermococcus sp. 2319x1                  EPLDRWILSRLHRLIKFATEELE-KYRFNLLTRELMTFVWHEVADDYIEMIKHRLYGEN-
Thermococcus litoralis                   EPLDRWILSRLHRLIKFATEELE-KYRFNLLTRELMTFVWHEVADDYIEMIKHCLYGED-
Thermococcales archaeon bit391           EPLDRWILSRLHRLIKFATEELE-KYRFNLLTRELMTFVWHEVADDYLEMIKHRLYGED-
Thermococcus sibiricus                   EPLDKWILSRLHRLIKFATEELE-KYRFNLLTRELMTFVWHEVADDYIEMIKHRLYGED-
Thermococci archaeon B45 G15             EPLDKWILSRLHRLINFATEELE-KYRFNLFTRELMNFVWHELADDYIEMIKHRLYGDD-
Palaeococcus pacificus                   EPLDRWILSRLHRLIKFATEEMD-SYRFNLFTRELMTFIWHEVADDYIEMIKYRLYGDD-
Thermococci archaeonB89 G9               EPIDRWILSRLHRLIRFATGEMD-SYRFNLFTRELMNFVWHEVADDYIEMIKYRLYGDD-
Thermococcus kodakarensis                EPLDRWILSRLHRLIKFATEEME-KYRFNLLTRELITFVWHEVADDYIEMIKYRLYGDD-
Pyrococcus furiosus                      EPLDRWILSRLHRIIKFATEELE-KYRFNLITRELMTFIWHEVADDYIEMIKHRLYGED-
Thermococci archaeon B61 G1              KPIDRWILYHLDQLIESTTNDFE-RYEFPQIMRRIWSFTWNKLADDYLEIVKHRIYDPKS
Thermococci archaeon B48 G16             KEIDRWILSKIDTLIEKVTLYLE-EYEYSRARTLIETSFWHDFCDNYLEIVKYRLYEKK-
Thermococci archaeon B54 G1              KEIDRWILSKIDTLIEKVTLYLE-EYEYSRARTLIETSFWHDFCDNYLEIVKYRLYEKK-
Euryarchaeota archaeon bit403            KIMDRWILSKYSKVVENATEYMD-EFRFDKAMKEIEEFLWHEFADHYIEIVKHRAYDND-
Mahella australiensis                    TLPDRWILTRYSQLVEEVTSNLE-RFELGLAAQKLYDFIWSEYCDWYIELSKVQLQQG--
Thermincola potens                       SPADKWIKSRFNKTVKTVTDALE-AYNLGEAAQSLYEFIWNEFCDWYIELVKPVLYGKE-
Thermofilum pendens                      SHADHWILRELASTVTRVTQALE-NYNFQEASQALVDFTWHKLADHYVEAVKHRLSRSD-
Thermoprotei archaeon bit383             TPIDHWILRELSSTIKEVTKSLE-EFDFQTASTKIIDFSWHKLCDHYLEAIKYRLSKED-
Crenarchaeota archaeon bit341            SLIDRWLLASLQELIQKVSEAFE-TFQFNTALEAIRNFTWHALADDYLEAVKHRLQTGRE
C.Verstraetearchaeota archaeon bit357    ELIDKWLLSKLEKLTFKVDEALS-KFQFNVALDAIRRFFWHVFCDHYIEAVKYRLYGGE-
C.Bathyarchaeota archaeon bit301         EPIDRWMLSKLERVKLKVTEALN-NYQFNIAVEALRGFIWHVLCDQYIEAVKYRLYNPEV
Hadesarchaea archaeon bit408             RLVDRWILSRFNRLIEQTTSWLE-DFQFNRALSAIQTFVWHELCDMYIEEVKHRLYGDD-
Methanothrix thermoacetophila            TQIDRWLLGELGMLVSKVTDAME-AFQFDEAFRAIRAFTWEVLADDYIEIVKSRLYGPD-
Methanoplanus limicola                   VLSDRWLMAKLSSAISEVTESMD-SYQYDRAVKTIREFAREIFADNYIEMVKGRLYGSG-
Geoglobus acetivorans                    RDADRWILSKLNRLIIEVDRSME-EYNFSNALKAIRSFTWYELADNYIEIVKNRLYSGS-
Ferroglobus placidus                     RDADRWILSKLNKLVKEVDEAME-NYSFSEALKLIRGFTWYEFADNYVELVKNRLYSGD-
Archaeoglobus fulgidus                   RDADRWILSKLNRLVGEVRKHMD-EYRFDEAIKAIRTFTWYEYADNYLEIVKNRLYSGS-
Halococcus saccharolyticus               TDADRWILSRCARVADEVAADMD-EYRFDRALRKVREFVWHDLADDYVELIKGRLYEGD-
Halalkalicoccus jeotgali                 RDADRWILTELDSVAEEVERDME-AYRFDAALRKIREFVWHDLADDYLELIKGRLYEGR-
Methanopyrus kandleri                    TPADRWILSKFHRLVDEVTEHLESGYRFNDAIKAIEEFAWEELADDYLEMAKLRLYRPEE
                                            *:*:          .   :. :        :        .* *:*  *
```

```
                                                  730       740       750       760       770       780
                                        .........|.........|.........|.........|.........|.........|
Thermococcus sp. 2319x1                 --EESKLKAKTALYELLYNITLLLAPFAPHITEELYQEMFRDKVGAKSVHLLSWPAYRED
Thermococcus litoralis                  --EESKLKAKAALYELLYNITLLLAPFAPHITEELYQEMFRDKVGAKSVHLLSWPAYRED
Thermococcales archaeon bit391          --EESKLKAKAALYELLYNITLLLAPFAPHITEELYQELFKDKVGAKSVHLLSWPTYRED
Thermococcus sibiricus                  --EESKLKAKMVLYELLYNITLLLGPLAPHITEELYQEMFKDKVGAKSVHLLEWPEYRED
Thermococci archaeon B45 G15            --EESKLKAKVALHELLYNIILLLAPLAPHITEELYQEMFKDKVGAKSLHLLEWPAYEED
Palaeococcus pacificus                  --EESKLKAKVALHELLYNILLLLAPLAPHITEELYQEVFKHKVGAKSIHLLEWPAYREE
Thermococci archaeonB89 G9              --EGSRLKAKVALHELLYNILLLLAPLAPHIAEEIYQDIFKPKIGAKSVHLLEWPKYTEE
Thermococcus kodakarensis               --EESKLKAKAALYELLYNVMLLLAPFVPHITEELYQNLFRERIGAKSVHLLEWPKYSEA
Pyrococcus furiosus                     --EESKLKAKVALYELLYNIMLLLAPFVPHITEELYHHIFKEKIGEKSVHLLQWPEYRED
Thermococci archaeon B61 G1             IEE--KRKTQFVLLKVLRDLVKLVAPITPFIAEEIYQEIDGR---KESVHIEEWPERIGS
Thermococci archaeon B48 G16            -----DRSAKYTVYNALLMYLKLFAIYIPHITEEIYQNIFKKFEEYDSIHISPWPEKLG-
Thermococci archaeon B54 G1             -----DRSAKYTVYNALLMYLKLFAIYIPHITEEIYQDIFKKFEEYDSIHISPWPEKLG-
Euryarchaeota archaeon bit403           ------ESAISTLYTVCSGVVKLMAPMMPHIAEEIYTMNFKDAEGDPSIHISSWPKPV--
Mahella australiensis                   --EAVKTNTLYVLCHVLDGILRLLHPYMPFITEEIWHHLPGS---YGTIMLADWPCVEAE
Thermincola potens                      -SPAARYTAQYVLWSVLEGTMRLLHPFMPFITEEIWQHLPHE---GKTIMTQQWPEFNTA
Thermofilum pendens                     ------EAAKYTLYVVLIKTLQMLSVFAPHISEEIYVDVLKKAGGWESITVSPWPEPP--
Thermoprotei archaeon bit383            ---KGSEAAKYTLYLVILRTLQMLSVFAPHVSEEIYHRLYEGKEKWKSITTSPWPSEE--
Crenarchaeota archaeon bit341           LSD--LTATQYCLREAVLTICKLLAPICPHISEAVYQQFPSE--ATKSVHLESWPEHEK-
C.Verstraetearchaeota archaeon bit357   --DESKRAAQYTLYEALFRILQLLAPFTPHIVEEIYQNIYAEDKGFLSIHKSSWPKVDEG
C.Bathyarchaeota archaeon bit301        YGEESRRAAQKTLYDVLYTSIQLLAPICPHITEEIYEALFKEHKGYVSIHVSPWPKPDRE
Hadesarchaea archaeon bit408            ---PTADAARYTLYHVMLTAAKLLAPFTPHFAEEIYHTYFAKDHPYPSVHASSWPEVNKK
Methanothrix thermoacetophila           --SDERRAAQATLYRVLDVLCRLMAPFIPFITEEIYTSLT-G---K-SVHTQSWPSLETY
Methanoplanus limicola                  --E-GKESALFALKTSVDALCRMLAPITPFFAEECYKHLTGG---KESVHERTWV--DYE
Geoglobus acetivorans                   --DEERVPAKYTLHRVLSTLIRLLAPITPFMAEECWQVFSGD---G-SVHLQRYPEADET
Ferroglobus placidus                    --EKEKLAARYTLYKVMNALIRLLAPFTPFLSEECWNVLFKD---G-SVHMQSYPEVEEK
Archaeoglobus fulgidus                  --EEEKRAAKFVLSYALDVLTRLLAPITPFMAEECWSHF-RE---G-SVHLQSYPVVEEE
Halococcus saccharolyticus              --PAERDAARHALSTTLSASLRLLAPFSPFLAEEAYHHLPDT---SGSVHAAAWPALDIA
Halalkalicoccus jeotgali                --PGERDAARHALYTALSGSLRMLSPFSPFFTEEVYRDLPGT---EGSVHTAEWPDVEAD
Methanopyrus kandleri                   LGEGSREAARAVLRHVLDGLLRLLAPFMPFVTEELYYRLFDE-----SVHDQAWPEASEK
                                                :   :         :.     *.. *  :           ::    :
```

```
                                              790       800       810       820       830       840
                                      .........|.........|.........|.........|.........|.........|
Thermococcus sp. 2319x1               RIDEEAERLGKLVSEIIGAMRRYKNSHGLALNAKLKHVAIYAID--SYEMLKALEKDIAG
Thermococcus litoralis                RIDEEAEKLGKFASEIIGAMRRYKNSHGLALNAKLKHVAIYATD--SYKMLKALEKDISG
Thermococcales archaeon bit391        RIDEEAEKLGKLASEIVGAMRRYKNSHGLALNAKLKHVAIYATD--SYDMLKAIEKDIAG
Thermococcus sibiricus                RVDEEAERIGKFASEIIGAMRRYKNSHGLALNAKLKHVAIYATD--SYEILKALERDIAG
Thermococci archaeon B45 G15          RIDEEAETLGKLASEIIGAMRKYKNSHGLALNAKLKHVAIYALD--SYDMLKALETDIAG
Palaeococcus pacificus                RVDEEAERLGEFASEVIGAMRRYKNSHGLSLNAKLKHVAIYTLD--SYEVLKQIEKDIAG
Thermococci archaeonB89 G9            RIDEEAEKLGELASEVVGIMRRYKNSHGLSLNTKLKHVAIYATD--SYDTFKSIEKDIAG
Thermococcus kodakarensis             RIDEEAEKLGELAREIVGAMRRYKNSHGLSLNAKLKHVAIYTTD--SYEVLKTIEKDIAG
Pyrococcus furiosus                   RIDEEAEKIGELAREIVSAMRKYKNSHGMPLNAKLKHVAIYATD--SYEMLKVIEKDIAG
Thermococci archaeon B61 G1           WLEEGA----EVSVLIIRGIRRYKGEKGIPLNRKLKEVEIFVNDQRLLGEIRRNLEDIEG
Thermococci archaeon B48 G16          ---IKGSNLGELCVSIISSLRKWKSDRGMALNAKIEDIVIFSSK-----DIDPIKEDIKR
Thermococci archaeon B54 G1           ---IKGSNLGELCVSIISSLRKWKSDRGMALNAKIEDIVIFSSK-----DIDPIKEDIKR
Euryarchaeota archaeon bit403         LTDEDAERKGELVKEIVSKIRGWKSEKGMSLSKEIDFVELVCEP----EKIIECKEDIAR
Mahella australiensis                 FVFEDSIDNMQSIMDVIQAIRNIKAEMNVEAGRKPKVVLVTDPS-Y-EPLFQANARYIER
Thermincola potens                    EVDEKAEAEMEMAMEVIKAVRNIRSEMNVAPSRKADVIINAGSG-AAMEVLNRCKPYLVN
Thermofilum pendens                   AYDEEKARVGDILIAVLAEGRRAKHDARIPLNKEVSAVYLYSEK--YSEELKAVVDDVAG
Thermoprotei archaeon bit383          EFNDKEAEIGDIAIAVIAEARRAKHDMKIALNANIAAIHIYSSK--YRDYLEKVAEDIKG
Crenarchaeota archaeon bit341         ALDDTSIRNGRFLLDLIAHARREKSSKGLSLGSNIKRIIVSTEA-EHLHLLKENEETILR
C.Verstraetearchaeota archaeon bit357 RINEKLEAVGDMIIAVIAELRRIKAERRIALSRPVEEAVIYASEDVKK-LLEEHLEIIGR
C.Bathyarchaeota archaeon bit301      RIDEEAEREGDLTIAVINAFRREKAGHRIPLNVKVDEATVYASSDWEAETVERNLEVIKG
Hadesarchaea archaeon bit408          FIDESAERAGELANLIVGALRQFKSERKMALSRKLPSVEIYASSKETAKQLKEVGADIVG
Methanothrix thermoacetophila         -----TSPEGALIREIAAAIRRYKSERGMALNAPLSGIEIYT--E--L---ELETFDLRG
Methanoplanus limicola                YSDSDAEVQGDLLVRVVSEVRRYKHDEGFALNAPLGHIVVYS--P--YE--INDGGDASS
Geoglobus acetivorans                 LIDENAERKGEMIKEIVEAIRRYKHDNGLALNAPMGRIGIFV--K---E--SIDTRDIAG
Ferroglobus placidus                  FIDEEAERRGEKIRKIVEEIRRYKHDRKMALNAPMKRILIYS--P---E--KFDVRDISG
Archaeoglobus fulgidus                FLDERAEKAGEEIKEIVAAVRKFKHDKGLALNAPLKKLIVYS--K--LD--GLDVRDIAG
Halococcus saccharolyticus            D--DAAEARGERIAAVASAVRAWKSDEGMALNADLDRIEVYA-DG--IA--ELDTGDLSA
Halalkalicoccus jeotgali              WDHEEATADGELIADAASTIRGWKSDSGMALNAELDRVELYAADD--IE--GLDTYDLSG
Methanopyrus kandleri                 WIDEGVEEVGEILREIVTEVRKAKTDAGLRMGAEFEGLTVHVQDEELAESLEKAIPDLKS
                                                          *  :    .  .
```

```
                                                    850       860       870       880       890       900
                                           .........|.........|.........|.........|.........|.........|
Thermococcus sp. 2319x1                    TMNIEKLEIIRGEP---ELEERITEIKPNFKTVGPKYGKLVPKIAAYLKE-N----AEEV
Thermococcus litoralis                     TMNIEKLEIIKGEP---ELEERIIEIKPNFKTVGPKYGKLVPKITAYLKE-N----AEEV
Thermococcales archaeon bit391             TMNIEKLEIIKGEP---ELEERITEIKPNFKTVGPKYGKLVPKITAYLKE-N----AEEV
Thermococcus sibiricus                     TMNIEKLEIIKGEP---ELEEKIAEIKPNFKTVGPKYGKLVPKITAYLKE-N----AEEV
Thermococci archaeon B45 G15               TMNIEKLEIIRGEP---DLEERIIEIKPNFKSVGPKYGRLVPKITAYLKE-N----ARKV
Palaeococcus pacificus                     TMNIEKLEVIKGEP---DLEERILEIKPNFKLVGPKYGKLVPKISAYLKE-N----AESV
Thermococci archaeonB89 G9                 TMNIEKLEIIKGEP---ELEERITEIKPNFKLVGPKYGKLVPKISKYLVE-N----AGEV
Thermococcus kodakarensis                  TMNIEKLEIIKGEP---ELEERIIEIKPNFKTVGPRYGKLVPKITAYLKE-N----AEEV
Pyrococcus furiosus                        TMNIERLEIVKGEP---QLEEKVVEIKPIYKRIGPRYGKLVPKIVKHLQE-N----AEEI
Thermococci archaeon B61 G1                ATRSEILI----REGVPEIDVVVKELEPDISKIGPEFKEKTKDVIEQMRKIKSEIGEEKL
Thermococci archaeon B48 G16               AMNVENLES---KKGKPEIEEKIVKVIPNYKVIGPIFGDRTKEIVKIIQDPE-------I
Thermococci archaeon B54 G1                AMNVENLEF---KKGKPEIEEKIVKVIPNYKVIGPIFGDRTKEIVKIIQDPE-------I
Euryarchaeota archaeon bit403              TVRAKELVV----AEKENLKENLVAVKPVYSKIGPVFKGKAEEIVEKLKTID----VGKL
Mahella australiensis                      LAGVSELSIKSDKSDISADAVS--AVTGRA-DIYIPLGELV----DI---------AKET
Thermincola potens                         LAHIANLTLEEKLDAEPEDAMS--AIIKGV-EIYMPLKGLV----DI---------QAEV
Thermofilum pendens                        TLRAKKVEVVRGEPA-GRKVP-----------EYPEISIQISP-----------------
Thermoprotei archaeon bit383               TIRAEKVIIHETGKG-ERQVP-----------EYPEITITLER-----------------
Crenarchaeota archaeon bit341              TLKADYMDLERLPLNSNQVGESS-----------AG--FVVEIIAK--------------
C.Verstraetearchaeota archaeon bit357      VIKARRVRVLSIAEGLGRTVR-----------DYPEIRVDIKV-----------------
C.Bathyarchaeota archaeon bit301           TCKIERIEV-YRSEGQGREVE-----------GYPNVKVALKV-----------------
Hadesarchaea archaeon bit408               TLRVGKLNI---KVGRPKLAERVLAVEPELAKLGPRLRADVKLIVQALRKAK----PEDL
Methanothrix thermoacetophila              VANAPIQ-L---RKGQPEIESRAVAVKPVMRFIGPRYKDQAGKIIKKLTSMD----PADV
Methanoplanus limicola                     TLNADVE-W---KCEKPELIKSVSDVRFNFGLIGPKFRKQANAYMNAIRALT----EDEK
Geoglobus acetivorans                      AVNAEVE-I---VEEMPEIQMGIAEVKPKFSIIGPKFRDRAGAVINAVKQMS----EAEI
Ferroglobus placidus                       AVNADVS-F---IEEMPKIETKVKEIKPKYSVIGPKFKEKAKKVVEAVTKLS----EEEK
Archaeoglobus fulgidus                     ATNSEVE-I---VTEMPEVRERVKELKPKFAIIGPMFREKAKTLIKAVGSLS----KEEK
Halococcus saccharolyticus                 TVNAPVT-V---ETGEPDVEMVAVGVDPDHSTIGPEFRDQAGAVVGALEAAD----PATI
Halalkalicoccus jeotgali                   AVNAPVY-I---ESGRPDVELVPVEIEPDQSVIGPEFRDRAGAVLGALSEAD----PAEI
Methanopyrus kandleri                      ATRAKEVEVEVGEP---KLERVPVKVEPRMDVIGPKYRELTRDIIEYVEN-N----PDEV
```

```
                                              910       920       930       940       950       960
                                     .........|.........|.........|.........|.........|.........|
Thermococcus sp. 2319x1              SRVLKEQGRVEFEV-------DGQKV---ELNKEDVV-IRKAV-FSEGEEVETAVVGDAV
Thermococcus litoralis               SRALKEQGKIEFEV-------DGQKV---ELNKEDIV-IRKAV-FSEGEEVETAVVGDAV
Thermococcales archaeon bit391       ARTLKEQGKVEFEV-------EGQKV---ELDKEDVV-IRKAV-FSEGEEVETAVVGDAV
Thermococcus sibiricus               AKALKEVGKVEFEV-------DGQKV---ELGKDDIV-LRKAV-FSEGEEVETAVVEDAV
Thermococci archaeon B45 G15         AKALKENGKVEFEA-------EEQKI---ELGKEDIV-IRKAV-FSEGEEVETVVVGDAV
Palaeococcus pacificus               AKALKESGKVEFEV-------EGGKV---ELGKEEIV-IRKAV-FSEGEEVKTAVVGDAV
Thermococci archaeonB89 G9           SIALKEKGKIEFEI-------DGQRI---TLGKDEVV-IRKTV-FSEGEEVETAVVGEAV
Thermococcus kodakarensis            AKALKESGKIEFEV-------DGQKV---ELTKDDIV-LRKAV-FSEGEEVETAVVGDAV
Pyrococcus furiosus                  GRRIKEEGKVEFEV-------EGQKV---VLEKEDIE-IKKAV-FSEGEEVETAVVRDAT
Thermococci archaeon B61 G1          YKSLLNSGIIELKL-------ESGET---VNVSE--------------------------
Thermococci archaeon B48 G16         AKK----------ID------SGEKVTEYKLCKDHISRIEKEY-RAEGRKVDILTGKDYI
Thermococci archaeon B54 G1          AKK----------ID------SGEKVTEYKLCKDHISRIEKEY-RAEGRKVDILTGKDYI
Euryarchaeota archaeon bit403        -----SEDEVNVRLD------SGETV---KLTKDFIN-FEKTV-TVKGKKVDVLNIGDTV
Mahella australiensis                ARLKKEYDELCAELDRTKAKLSNQSFI--TKAPTAVVQAERDK------AVQYQAMIDSI
Thermincola potens                   ARLRKELDVLEKELARVNGKLNNQKFL--DKAPEDVIAKEKAK------HAEYSEKKQAV
Thermofilum pendens                  ------------------------------------------------------------
Thermoprotei archaeon bit383         ------------------------------------------------------------
Crenarchaeota archaeon bit341        ------------------------------------------------------------
C.Verstraetearchaeota archaeon bit357 ------------------------------------------------------------
C.Bathyarchaeota archaeon bit301     ------------------------------------------------------------
Hadesarchaea archaeon bit408         AK-QLAQGKLKLRA-------GGKRF---ELAPEDVR-VIKET-ARAGRKVEIIDIQKPA
Methanothrix thermoacetophila        ERML-ASGRVEIEG-------A-------EITPEMVE-IVRET-LSMGEAVDVLRLDRAT
Methanoplanus limicola               ITP-----PRTVIM-------DGVET---EVLENSFE-PVYSY-SVAGEKVDLLTLTDDV
Geoglobus acetivorans                LKFI-NEGRVTLSI-------NGEEY---ELESEWFE-FKFEK-SISGKNVDVVETSSAV
Ferroglobus placidus                 FKFM-ETGKLEVEV-------NGERI---ELSSDWFE-FKLEK-VVGDLSVDVLEVENVV
Archaeoglobus fulgidus               ERLL-KEGAIQVNL-------DGASV---EVKAEWFE-AVTEK-SIEGREVEMLETANSV
Halococcus saccharolyticus           ERQKREDGRIEIDL-------DGETV---TLDGDAVA-VEREQR-VAGEEVAVLDAEDTT
Halalkalicoccus jeotgali             KAQK-QTGEIELDI-------GEETV---ALSPEAVS-VHEERRAESGEEVAVLETERAT
Methanopyrus kandleri                ASAIKEDGKAKLEI-------DGEKV---VLDEECVD-VEWEL-RVKGGEGKAVEIRPGV
```

```
                                                      970       980       990
                                             .........|.........|.........|
Thermococcus sp. 2319x1                      ILFF--------------------
Thermococcus litoralis                       ILFF--------------------
Thermococcales archaeon bit391               ILFF--------------------
Thermococcus sibiricus                       IVFF--------------------
Thermococci archaeon B45 G15                 ILFF--------------------
Palaeococcus pacificus                       ILFFS-------------------
Thermococci archaeonB89 G9                   ILFFA-------------------
Thermococcus kodakarensis                    ILFF--------------------
Pyrococcus furiosus                          ILFF--------------------
Thermococci archaeon B61 G1                  ------------------------
Thermococci archaeon B48 G16                 IEIF--------------------
Thermococci archaeon B54 G1                  IEIF--------------------
Euryarchaeota archaeon bit403                ALIGRV------------------
Mahella australiensis                        KERLEHLEK---------------
Thermincola potens                           LERLAKLGVK--------------
Thermofilum pendens                          ------------------------
Thermoprotei archaeon bit383                 ------------------------
Crenarchaeota archaeon bit341                ------------------------
C.Verstraetearchaeota archaeon bit357        ------------------------
C.Bathyarchaeota archaeon bit301             ------------------------
Hadesarchaea archaeon bit408                 LTLLITVPRRSLVRRRTRSARKAR
Methanothrix thermoacetophila                LLIRRSS-----------------
Methanoplanus limicola                       ILTVQKKE----------------
Geoglobus acetivorans                        IIIEKR------------------
Ferroglobus placidus                         VMIERV------------------
Archaeoglobus fulgidus                       VFVEV-------------------
Halococcus saccharolyticus                   VLVFP-------------------
Halalkalicoccus jeotgali                     VLVFP-------------------
Methanopyrus kandleri                        VVEI-----RGLST----------
```
