## Supplementary Table 1 and 2 for "*Thermococci*-to-*Clostridia* Pathway for the Evolution of the Bacteria Domain"

**Supplementary Table 1. Bacterial species with relatively high VARS-IARS bitscores (ref. 9).**

| Species name | VARS-IARS Bitscore | GenBank VARS Accession | GenBank IARS Accession |
|---|---|---|---|
| *Mahella australiensis 50-1 BON* | 378 | AEE96871.1 | AEE96541.1 |
| *Thermincola potens JR* | 377 | ADG83394.1 | ADG82707.1 |
| *Halobacteroides halobius DSM 5150* | 375 | AGB41920.1 | AGB40823.1 |
| *Halothermothrix orenii H 168* | 373 | ACL70208.1 | ACL69683.1 |
| *Desulfosporosinus orientis DSM 765* | 372 | AET70613.1 | AET70197.1 |
| *Caldicellulosiruptor lactoaceticus 6A* | 368 | AEM73516.1 | AEM73389.1 |
| *Carboxydocella thermautotrophica* | 366 | AVX31974.1 | AVX31394.1 |
| *Thermotoga sp. RQ7* | 363 | AJG40845.1 | AJG41316.1 |
| *Moorella thermoacetica ATCC 39073* | 361 | ABC18862.1 | ABC19180.1 |
| *Marinitoga sp. 1137* | 355 | APT76568.1 | APT76656.1 |
| *Caldanaerobacter subterraneus subsp. tengcongensis MB4* | 353 | AAM24038.1 | AAM24798.1 |
| *Carboxydothermus hydrogenoformans Z-2901* | 353 | ABB15424.1 | ABB14741.1 |
| *Pelotomaculum thermopropionicum SI* | 353 | BAF58993.1 | BAF60001.1 |
| *Thermosipho africanus TCF52B* | 352 | ACJ76327.1 | ACJ75295.1 |
| *Fervidobacterium pennivorans DSM 9078* | 350 | AFG34874.1 | AFG35844.1 |
| *Thermoanaerobacter mathranii subsp. mathranii str. A3* | 349 | ADH60525.1 | ADH61144.1 |
| *Desulfitobacterium dehalogenans ATCC 51507* | 347 | AFM02008.1 | AFM01700.1 |
| *Acidaminococcus fermentans DSM 20731* | 347 | ADB46800.1 | ADB47854.1 |
| *Thermoclostridium stercorarium* | 345 | AGC69102.1 | AGC67610.1 |
| *Pseudothermotoga lettingae TMO* | 343 | ABV34537.1 | ABV32949.1 |
| *Desulfotomaculum ferrireducens* | 343 | AQS59968.1 | AQS58345.1 |
| *Hungateiclostridium thermocellum ATCC 27405* | 342 | ABN51562.1 | ABN52022.1 |
| *Kosmotoga olearia TBF 19.5.1* | 340 | ACR80423.1 | ACR78740.1 |
| *Halanaerobium hydrogeniformans* | 340 | ADQ14528.1 | ADQ15019.1 |
| *Anoxybacter fermentans* | 339 | AZR73355.1 | AZR74642.1 |
| *Aneurinibacillus sp. XH2* | 337 | AMA72103.1 | AMA72590.1 |
| *Syntrophobotulus glycolicus DSM 8271* | 337 | ADY56765.1 | ADY55025.1 |
| *Thermoanaerobacterium thermosaccharolyticum DSM 571* | 335 | ADL68565.1 | ADL68740.1 |
| *Desulfofarcimen acetoxidans DSM 771* | 332 | ACV64078.1 | ACV61975.1 |
| *Clostridium sp. BNL1100* | 332 | AEY67008.1 | AEY66369.1 |
| *Desulfallas gibsoniae DSM 7213* | 332 | AGL02659.1 | AGL02937.1 |
| *Thermodesulfatator indicus DSM 15286* | 330 | AEH44784.1 | AEH45247.1 |
| *Syntrophothermus lipocalidus DSM 12680* | 330 | ADI01463.1 | ADI01607.1 |
| *Ruminiclostridium cellulolyticum H10* | 330 | ACL75556.1 | ACL76203.1 |
| *Novibacillus thermophilus* | 330 | AQS55176.1 | AQS55533.1 |
| *Pelosinus sp. UFO1* | 330 | AIF52510.1 | AIF51959.1 |
| *Thermosediminibacter oceani DSM 16646* | 328 | ADL07455.1 | ADL08379.1 |
| *Natranaerobius thermophilus JW/NM-WN-LF* | 327 | ACB84692.1 | ACB84903.1 |
| *Petrotoga mobilis SJ95* | 326 | ABX32160.1 | ABX32320.1 |
| *Desulfofundulus kuznetsovii DSM 6115* | 326 | AEG14027.1 | AEG14774.1 |
| *Dictyoglomus turgidum DSM 6724* | 325 | ACK42369.1 | ACK42703.1 |
| *Thermaerobacter marianensis DSM 12885* | 325 | ADU52301.1 | ADU50999.1 |
| *Thermacetogenium phaeum DSM 12270* | 324 | AFV10733.1 | AFV11511.1 |
| *Candidatus Desulforudis audaxviator MP104C* | 323 | ACA59979.1 | ACA59917.1 |
| *Desulfurobacterium thermolithotrophum DSM 11699* | 323 | ADY74095.1 | ADY73289.1 |

| | | | |
|---|---|---|---|
| *Thermovibrio ammonificans HB-1* | 322 | ADU97632.1 | ADU96612.1 |
| *Defluviitoga tunisiensis* | 321 | CEP78993.1 | CEP79118.1 |
| *Acetohalobium arabaticum DSM 5501* | 320 | ADL12058.1 | ADL12417.1 |
| *Syntrophomonas wolfei subsp. wolfei str. Goettingen G311* | 319 | ABI68948.1 | ABI68277.1 |
| *Ammonifex degensii KC4* | 319 | ACX51457.1 | ACX52153.1 |
| *Cloacibacillus porcorum* | 319 | ANZ45035.1 | ANZ45058.1 |
| *Thermoactinomyces vulgaris* | 318 | QBK14449.1 | QBK13828.1 |
| *Thermodesulfobacterium geofontis OPF15* | 318 | AEH23131.1 | AEH23000.1 |
| *Megasphaera stantonii* | 317 | AXL21685.1 | AXL20633.1 |
| *Eubacterium limosum* | 317 | ARD67998.1 | ARD64226.1 |
| *Paenibacillus swuensis* | 314 | ANE47159.1 | ANE47386.1 |
| *Pseudoclostridium thermosuccinogenes* | 314 | AUS97845.1 | AUS96188.1 |
| *Christensenella minuta* | 313 | AYH39911.1 | AYH41262.1 |
| *Megamonas hypermegale* | 313 | SNV00186.1 | SNU99632.1 |
| *Caproiciproducens sp. NJN-50* | 312 | QAT50132.1 | QAT49841.1 |
| *Veillonella rodentium* | 312 | SNV70071.1 | SNV56080.1 |
| *Fusobacterium ulcerans* | 312 | AVQ26760.1 | AVQ28042.1 |
| *Halobacillus halophilus DSM 2266* | 311 | CCG46018.1 | CCG43613.1 |
| *Kyrpidia spormannii* | 310 | ATY84344.1 | ATY84742.1 |
| *Fastidiosipila sanguinis* | 310 | AVM42064.1 | AVM42330.1 |
| *Geobacter anodireducens* | 309 | ANA40045.1 | ANA41297.1 |
| *Laceyella sacchari* | 308 | AUS08415.1 | AUS09329.1 |
| *Syntrophus aciditrophicus SB* | 308 | ABC77822.1 | ABC77375.1 |
| *Thermosulfidibacter takaii ABI70S6* | 308 | BAT71800.1 | BAT71422.1 |
| *Geobacillus sp. Y4.1MC1* | 308 | ADP73746.1 | ADP75454.1 |
| *Parageobacillus thermoglucosidasius* | 308 | ALF11693.1 | ALF10077.1 |
| *Caldimicrobium thiodismutans* | 308 | BAU23034.1 | BAU23965.1 |
| *Heliobacterium modesticaldum Ice1* | 308 | ABZ85278.1 | ABZ84653.1 |
| *Brevibacillus brevis NBRC 100599* | 308 | BAH42801.1 | BAH44795.1 |
| *Acetobacterium woodii DSM 1030* | 307 | AFA48747.1 | AFA47645.1 |
| *Ruminococcus bicirculans* | 306 | CCO04152.1 | CCO04153.1 |
| *Desulfurella acetivorans A63* | 305 | AHF96597.1 | AHF96772.1 |
| *Intestinimonas butyriciproducens* | 305 | ALP93554.1 | ALP92579.1 |
| *Dialister pneumosintes* | 304 | AOH38878.1 | AOH39427.1 |
| *Alkaliphilus oremlandii OhILAs* | 304 | ABW19574.1 | ABW19732.1 |
| *Desulfobacca acetoxidans DSM 11109* | 303 | AEB09271.1 | AEB10708.1 |
| *Cryptobacterium curtum DSM 15641* | 303 | ACU94599.1 | ACU94350.1 |
| *Pelobacter propionicus DSM 2379* | 302 | ABK99195.1 | ABL01188.1 |
| *Kurthia sp. 11kri321* | 302 | AMA61894.1 | AMA61845.1 |
| *Methylomusa anaerophila* | 302 | BBB89773.1 | BBB93010.1 |
| *[Clostridium] saccharolyticum WM1* | 302 | ADL04925.1 | ADL05433.1 |
| *Thermobacillus composti KWC4* | 302 | AGA58663.1 | AGA58411.1 |
| *Geosporobacter ferrireducens* | 302 | AOT68849.1 | AOT72150.1 |
| *Sphaerobacter thermophilus DSM 20745* | 301 | ACZ39046.1 | ACZ37776.1 |
| *Ethanoligenens harbinense YUAN-3* | 300 | ADU27025.1 | ADU26270.1 |
| *[Bacillus] caldolyticus* | 299 | AUI37651.1 | AUI35899.1 |
| *Aminobacterium colombiense DSM 12261* | 298 | ADE57025.1 | ADE57086.1 |
| *Bacillus smithii* | 298 | AKP47994.1 | AKP46785.1 |
| *Geoalkalibacter subterraneus* | 298 | AJF07012.1 | AJF05548.1 |

| | | | |
|---|---|---|---|
| *Symbiobacterium thermophilum IAM 14863* | 298 | BAD39351.1 | BAD40216.1 |
| *Slackia heliotrinireducens DSM 20476* | 298 | ACV22207.1 | ACV22123.1 |
| *Anoxybacillus amylolyticus* | 296 | ANB60387.1 | ANB59918.1 |
| *Thermanaerovibrio acidaminovorans DSM 6589* | 296 | ACZ19182.1 | ACZ19217.1 |
| *Oscillibacter valericigenes Sjm18-20* | 296 | BAK98761.1 | BAL01799.1 |
| *Desulfurivibrio alkaliphilus AHT 2* | 296 | ADH85749.1 | ADH85876.1 |
| *Tepidanaerobacter acetatoxydans Re1* | 296 | CCP26388.1 | CCP25437.1 |
| *Acholeplasma palmae J233* | 295 | CCV64478.1 | CCV64446.1 |
| *Carnobacterium sp. CP1* | 295 | ALV20826.1 | ALV20798.1 |
| *Desulfuromonas sp. DDH964* | 295 | AMV72963.1 | AMV71142.1 |
| *Anaerostipes hadrus* | 295 | AQP40389.1 | AQP39491.1 |
| *Rummeliibacillus stabekisii* | 293 | AMW99513.1 | AMW98354.1 |
| *Aerococcus urinae* | 293 | AMB96652.1 | AMB95533.1 |
| *Fictibacillus phosphorivorans* | 293 | ANC78039.1 | ANC76790.1 |
| *Syntrophobacter fumaroxidans MPOB* | 293 | ABK15894.1 | ABK15825.1 |
| *Phascolarctobacterium faecium* | 293 | BBG63265.1 | BBG63792.1 |
| *Blautia hansenii DSM 20583* | 293 | ASM70319.1 | ASM70670.1 |
| *Selenomonas sputigena ATCC 35185* | 292 | AEB99470.1 | AEC00247.1 |
| *Mesotoga infera* | 291 | SSC13855.1 | SSC11579.1 |
| *Thermomicrobium roseum DSM 5159* | 291 | ACM06199.1 | ACM05907.1 |
| *Thermovirga lienii DSM 17291* | 290 | AER66797.1 | AER66666.1 |
| *Desulfotalea psychrophila LSv54* | 290 | CAG36525.1 | CAG37281.1 |
| *Desulfatibacillum alkenivorans AK-01* | 288 | ACL02204.1 | ACL03467.1 |
| *Desulfomonile tiedjei DSM 6799* | 288 | AFM26329.1 | AFM26721.1 |
| *[Eubacterium] eligens ATCC 27750* | 288 | ACR72289.1 | ACR71076.1 |
| *Enterococcus faecalis V583* | 287 | AAO82619.1 | AAO80809.1 |
| *Lysinibacillus sp. 2017* | 287 | AWE08182.1 | AWE07832.1 |
| *Tumebacillus algifaecis* | 286 | ASS74933.1 | ASS75836.1 |
| *Turicibacter sp. H121* | 286 | AMC07945.1 | AMC07628.1 |
| *Eggerthella sp. YY7918* | 286 | BAK45005.1 | BAK44631.1 |
| *Thermobaculum terrenum ATCC BAA-798* | 286 | ACZ42389.1 | ACZ41894.1 |
| *Leptotrichia sp. oral taxon 498* | 285 | ASQ49265.1 | ASQ49337.1 |
| *Oscillatoria acuminata PCC 6304* | 285 | AFY79923.1 | AFY83471.1 |
| *Solibacillus silvestris* | 284 | AMO86189.1 | AMO84606.1 |
| *Desulfohalobium retbaense DSM 5692* | 284 | ACV67553.1 | ACV68406.1 |
| *Limnochorda pilosa* | 283 | BAS28631.1 | BAS27387.1 |
| *Peptoniphilus harei* | 283 | VEJ34856.1 | VEJ33939.1 |
| *Acetoanaerobium sticklandii* | 283 | CBH20748.1 | CBH21996.1 |
| *Dehalobacter restrictus DSM 9455* | 283 | AHF10837.1 | AHF09194.1 |
| *Desulfarculus baarsii DSM 2075* | 282 | ADK85366.1 | ADK83788.1 |
| *Butyrivibrio hungatei* | 282 | AOZ95720.1 | AOZ97709.1 |
| *Virgibacillus necropolis* | 281 | ASN05094.1 | ASN05837.1 |
| *Aeribacillus pallidus* | 281 | ASS92063.1 | ASS91633.1 |
| *Pseudodesulfovibrio indicus* | 281 | AMK10483.1 | AMK11866.1 |
| *Vampirococcus sp. LiM* | 281 | QAT17095.1 | QAT17258.1 |
| *Thermodesulfobium acidiphilum* | 280 | AWB10652.1 | AWB10402.1 |
| *Flavonifractor plautii* | 280 | ANU41938.1 | ANU42798.1 |
| *Flexistipes sinusarabici DSM 4947* | 280 | AEI14666.1 | AEI14462.1 |
| *Paeniclostridium sordellii* | 280 | AUN13047.1 | AUN15436.1 |

| | | | |
|---|---|---|---|
| *Cohnella sp. 18JY8-7* | 280 | AYQ73220.1 | AYQ71818.1 |
| *Gottschalkia acidurici 9a* | 280 | AFS77895.1 | AFS77688.1 |
| *Desulfococcus multivorans* | 279 | AOY60649.1 | AOY58831.1 |
| *Calditerrivibrio nitroreducens DSM 19672* | 278 | ADR19219.1 | ADR18993.1 |
| *Thermus oshimai JL-2* | 278 | AFV76228.1 | AFV76495.1 |
| *Halocella sp. SP3-1* | 278 | AZO94277.1 | AZO95111.1 |
| *Alicyclobacillus acidocaldarius* | 278 | ACV58843.1 | ACV59381.1 |
| *Cellulosilyticum lentocellum DSM 5427* | 278 | ADZ83647.1 | ADZ85330.1 |
| *Caldisericum exile AZM16c01* | 278 | BAL81338.1 | BAL80506.1 |
| *Desulfobacterium autotrophicum HRM2* | 277 | ACN15188.1 | ACN15583.1 |
| *Desulfomicrobium orale DSM 12838* | 277 | AMD93679.1 | AMD93152.1 |
| *Kitasatospora sp. MMS16-BH015* | 276 | AUG77222.1 | AUG76718.1 |
| *Clostridioides difficile 630* | 276 | CAJ70153.1 | CAJ69504.1 |
| *Jeotgalibacillus malaysiensis* | 275 | AJD91673.1 | AJD90928.1 |
| *Paenisporosarcina sp. K2R23-3* | 275 | AYC29766.1 | AYC29176.1 |
| *Lactobacillus ginsenosidimutans* | 275 | AKP66397.1 | AKP66419.1 |
| *Tetragenococcus koreensis* | 275 | AYW45599.1 | AYW44537.1 |
| *Sporanaerobacter sp. NJN-17* | 275 | QAT61311.1 | QAT62717.1 |
| *Treponema caldarium DSM 7334* | 275 | AEJ19967.1 | AEJ19327.1 |
| *Oceanobacillus iheyensis HTE831* | 275 | BAC14017.1 | BAC12318.1 |
| *Vagococcus teuberi* | 274 | APB31984.1 | APB30842.1 |
| *Finegoldia magna ATCC 29328* | 274 | BAG08225.1 | BAG08522.1 |
| *Denitrobacterium detoxificans* | 273 | ANE22847.1 | ANE22797.1 |
| *Acetomicrobium mobile DSM 13181* | 273 | AFM21392.1 | AFM21556.1 |
| *Melissococcus plutonius S1* | 273 | AIM24421.1 | AIM24781.1 |
| *Listeria seeligeri serovar 1/2b str. SLCC3954* | 272 | CBH27618.1 | CBH28152.1 |
| *Adlercreutzia equolifaciens DSM 19450* | 272 | BAN77119.1 | BAN76975.1 |
| *Thermodesulfovibrio yellowstonii DSM 11347* | 272 | ACI20807.1 | ACI21479.1 |
| *Hippea maritima DSM 10411* | 271 | AEA33982.1 | AEA33278.1 |
| *Lactococcus raffinolactis* | 271 | ATC61001.1 | ATC60925.1 |
| *Salimicrobium jeotgali* | 271 | AKG04207.1 | AKG05435.1 |
| *Lachnoanaerobaculum umeaense* | 271 | AYA99829.1 | AYA99299.1 |
| *Coprothermobacter proteolyticus DSM 5265* | 270 | ACI16878.1 | ACI17812.1 |
| *Desulfobulbus propionicus DSM 2032* | 270 | ADW19170.1 | ADW18535.1 |
| *Coriobacterium glomerans PW2* | 270 | AEB06921.1 | AEB07126.1 |
| *Egibacter rhizosphaerae* | 270 | QBI18667.1 | QBI18122.1 |
| *Streptobacillus moniliformis DSM 12112* | 269 | ACZ00721.1 | ACZ01556.1 |
| *Deferribacter desulfuricans SSM1* | 269 | BAI80001.1 | BAI80654.1 |
| *Synechococcus sp. RCC307* | 269 | CAK29319.1 | CAK29115.1 |
| *Geitlerinema sp. PCC 7407* | 269 | AFY65874.1 | AFY67335.1 |
| *Endomicrobium proavitum* | 268 | AKL97525.1 | AKL97543.1 |
| *Brachyspira murdochii DSM 12563* | 268 | ADG72641.1 | ADG71970.1 |
| *Marinilactibacillus sp. 15R* | 268 | API89175.1 | API89156.1 |
| *Sulfobacillus acidophilus TPY* | 268 | AEJ38797.1 | AEJ41607.1 |
| *Lachnoclostridium sp. YL32* | 268 | ANU50116.1 | ANU45319.1 |
| *Elusimicrobium minutum Pei191* | 267 | ACC97815.1 | ACC98696.1 |
| *Dehalogenimonas lykanthroporepellens BL-DC-9* | 267 | ADJ26336.1 | ADJ26052.1 |
| *Streptacidiphilus sp. DSM 106435* | 267 | AXI77561.1 | AXI77076.1 |
| *Peptoclostridium acidaminophilum DSM 3953* | 267 | AHM56900.1 | AHM56942.1 |

| | | | |
|---|---|---|---|
| *Sulfurihydrogenibium azorense Az-Fu1* | 267 | ACN99343.1 | ACN98511.1 |
| *Microcoleus sp. PCC 7113* | 267 | AFZ16060.1 | AFZ19256.1 |
| *Desulfobacula toluolica Tol2* | 266 | CCK79813.1 | CCK80266.1 |
| *Calothrix sp. NIES-3974* | 266 | BAZ05233.1 | BAZ05421.1 |
| *Staphylococcus lutrae* | 265 | ARJ49959.1 | ARJ51623.1 |
| *Desulfocapsa sulfexigens DSM 10523* | 265 | AGF76766.1 | AGF79927.1 |
| *Leptolyngbya sp. NIES-3755* | 265 | BAU11788.1 | BAU15001.1 |
| *Anaeromyxobacter dehalogenans 2CP-1* | 265 | ACL64913.1 | ACL63373.1 |
| *Desulfovibrio fairfieldensis* | 264 | AMD88656.1 | AMD88804.1 |
| *Spirochaeta thermophila DSM 6578* | 264 | AEJ60725.1 | AEJ61905.1 |
| *Streptomyces xiamenensis* | 262 | AKG42943.1 | AKG42400.1 |
| *Amphibacillus xylanus NBRC 15112* | 261 | BAM47129.1 | BAM47700.1 |
| *Faecalibacterium prausnitzii* | 261 | AXB29292.1 | AXB28767.1 |
| *Sediminispirochaeta smaragdinae DSM 11293* | 261 | ADK81267.1 | ADK81020.1 |
| *Nostoc punctiforme PCC 73102* | 261 | ACC83038.1 | ACC80175.1 |
| *Brochothrix thermosphacta* | 260 | ATF27190.1 | ATF25284.1 |
| *Geovibrio thiophilus* | 259 | QAR33016.1 | QAR32883.1 |
| *Gemella haemolysans* | 259 | VEI39080.1 | VEI38737.1 |
| *Sebaldella termitidis ATCC 33386* | 259 | ACZ08487.1 | ACZ10771.1 |
| *Chondrocystis sp. NIES-4102* | 259 | BAZ44833.1 | BAZ46643.1 |
| *Nitrospira japonica* | 259 | SLM49140.1 | SLM48255.1 |
| *Chroococcidiopsis thermalis PCC 7203* | 259 | AFY86310.1 | AFY86358.1 |
| *Gloeocapsa sp. PCC 7428* | 259 | AFZ31327.1 | AFZ28738.1 |
| *Nodularia spumigena UHCC 0039* | 259 | AVZ29549.1 | AVZ30146.1 |
| *Melittangium boletus DSM 14713* | 259 | ATB34408.1 | ATB30341.1 |
| *Sporolactobacillus terrae* | 258 | QAA22933.1 | QAA22279.1 |
| *Macrococcus canis* | 258 | ARQ07333.1 | ARQ06641.1 |
| *Planococcus sp. PAMC 21323* | 258 | AIY05145.1 | AIY05925.1 |
| *Archangium gephyra* | 258 | AKJ02821.1 | AKI99538.1 |
| *Filifactor alocis ATCC 35896* | 257 | EFE28118.1 | EFE27741.1 |
| *Dactylococcopsis salina PCC 8305* | 257 | AFZ50261.1 | AFZ50190.1 |
| *[Brevibacterium] frigoritolerans* | 255 | AZV63346.1 | AZV60600.1 |
| *Parolsenella catena* | 254 | BBH50018.1 | BBH50120.1 |
| *Anaerotignum propionicum DSM 1682* | 254 | AMJ40794.1 | AMJ40272.1 |
| *Actinoplanes sp. N902-109* | 254 | AGL15016.1 | AGL18846.1 |
| *Jeotgalibaca dankookensis* | 253 | AQS52858.1 | AQS52845.1 |
| *Sporosarcina sp. P37* | 253 | ARK23626.1 | ARK25708.1 |
| *Actinoalloteichus sp. AHMU CJ021* | 253 | AUS80007.1 | AUS79738.1 |
| *Myxococcus xanthus DK 1622* | 253 | ABF86989.1 | ABF86287.1 |
| *Corallococcus coralloides DSM 2259* | 252 | AFE10628.1 | AFE09121.1 |
| *Exiguobacterium sp. AT1b* | 251 | ACQ71563.1 | ACQ71761.1 |
| *Terribacillus goriensis* | 251 | AIF66976.1 | AIF66300.1 |
| *Weissella koreensis KACC 15510* | 251 | AEJ23200.1 | AEJ23114.1 |
| *Synechocystis sp. IPPAS B-1465* | 251 | AVP90913.1 | AVP89093.1 |
| *Salinispira pacifica* | 251 | AHC14211.1 | AHC14223.1 |
| *Dehalococcoides sp. UCH007* | 250 | BAQ34320.1 | BAQ34830.1 |
| *Ilyobacter polytropus DSM 2926* | 249 | ADO82367.1 | ADO82448.1 |
| *Oceanithermus profundus DSM 14977* | 249 | ADR36799.1 | ADR36604.1 |
| *Parvimonas micra* | 249 | AIZ36585.1 | AIZ36140.1 |

| | | | |
|---|---|---|---|
| *Trichormus variabilis ATCC 29413* | 249 | ABA22592.1 | ABA23379.1 |
| *Fischerella sp. NIES-3754* | 249 | BAU04776.1 | BAU08255.1 |
| *Thermosynechococcus elongatus BP-1* | 248 | BAC09055.1 | BAC09882.1 |
| *Rubrobacter xylanophilus DSM 9941* | 248 | ABG04497.1 | ABG05355.1 |
| *Bdellovibrio bacteriovorus HD100* | 248 | CAE78439.1 | CAE80087.1 |
| *Pseudanabaena sp. ABRG5-3* | 247 | BBC24721.1 | BBC26310.1 |
| *Crinalium epipsammum PCC 9333* | 247 | AFZ11058.1 | AFZ13052.1 |
| *Thermobispora bispora DSM 43833* | 246 | ADG89177.1 | ADG88134.1 |
| *Cyanothece sp. PCC 7425* | 246 | ACL43456.1 | ACL45442.1 |

**Supplementary Table 2. Archaeal species with relatively high VARS-IARS bitscores. See ref. 43 for sequences of *C.Hydrothermarchaeota archaeon_B51_G15*, *Thermococci archaeon_B45_G15, Thermococci archaeon_B89_G9, Thermococci archaeon_B61_G1*, *Thermococci archaeon_B88_G9*, *Thermococci archaeon_B48_G16*, *Thermococci archaeon_B54_G1* and *Thermoprotei archaeon_B75_G16*; ref. 46 for sequence of *Euryarchaeota archaeon_bit403*; and ref. 9 for remaining entries.**

| Species name | VARS-IARS Bitscore | GenBank VARS Accession | GenBank IARS Accession |
|---|---|---|---|
| *Methanothermobacter tenebrarum* | 466 | RAO79057.1 | RAO78531.1 |
| *Ferroglobus placidus DSM 10642* | 462 | ADC64350.1 | ADC65758.1 |
| *Methanothermobacter sp. EMTCatA1* | 462 | BAZ98807.1 | BAZ99378.1 |
| *Methanothermobacter sp. CaT2* | 461 | BAM69935.1 | BAM70497.1 |
| *Methanothermobacter wolfeii* | 460 | SCM57570.1 | SCM58638.1 |
| *Methanothermobacter thermautotrophicus str. Delta H* | 459 | AAB85270.1 | AAB85852.1 |
| *Methanothermobacter defluvii* | 458 | REE28596.1 | REE28027.1 |
| *C.Syntrophoarchaeum butanivorans* | 451 | OFV66237.1 | OFV66061.1 |
| *Methanothermobacter marburgensis str. Marburg* | 451 | ADL58755.1 | ADL59330.1 |
| *Geoglobus ahangari* | 449 | AKG90833.1 | AKG91282.1 |
| *Methanothermobacter sp. MT-2* | 448 | BAW31196.1 | BAW30730.1 |
| *Methanobacteriaceae archaeon 41_258* | 444 | KUK00435.1 | KUK01125.1 |
| *C.Hydrothermarchaeota archaeon_B51_G15* | 439 | RLG58128.1 | RLG56379.1 |
| *Methanothermus fervidus DSM 2088* | 436 | ADP77172.1 | ADP77910.1 |
| *Hadesarchaea archaeon YNP_45* | 425 | KUO39405.1 | KUO39832.1 |
| *Archaeoglobales archaeon* | 421 | RLI83164.1 | RLI83421.1 |
| *Geoglobus acetivorans* | 418 | AIY90211.1 | AIY90260.1 |
| *Archaeoglobus profundus DSM 5631* | 416 | ADB57081.1 | ADB57431.1 |
| *Archaeoglobus sulfaticallidus PM70-1* | 412 | AGK60696.1 | AGK60827.1 |
| *Hadesarchaea archaeon_bit408 DG-33-1* | 408 | KUO40919.1 | KUO41495.1 |
| *Archaeoglobales archaeon* | 406 | RLI80090.1 | RLI78156.1 |
| *Euryarchaeota archaeon_bit405 55_53* | 405 | KUK04811.1 | KUK03924.1 |
| *Methanosarcinales archeaon 56_1174* | 405 | KUK30140.1 | KUK29494.1 |
| *Methanobacterium sp. BRmetb2* | 404 | AXV36984.1 | AXV37641.1 |
| *Methanohalophilus halophilus* | 403 | APH38319.1 | APH39716.1 |
| *Methanobacterium sp. A39* | 403 | OEC86700.1 | OEC86557.1 |
| *Euryarchaeota archaeon_bit403 CG_4_9_14_3_um_filter_38_12* | 403 | PJB21536.1 | PJB21216.1 |
| *Methanohalophilus sp. WG1-DM* | 402 | RXG33700.1 | RXG35309.1 |
| *Methanohalophilus mahii DSM 5219* | 402 | ADE35899.1 | ADE36306.1 |
| *Methanobacterium sp.* | 402 | RJS48095.1 | RJS49886.1 |
| *Methanobacterium congolense* | 402 | SCG85805.1 | SCG86792.1 |
| *Methanobacterium bryantii* | 402 | PAV03978.1 | PAV03078.1 |
| *Methanohalophilus sp. 2-GBenrich* | 401 | ODV49694.1 | ODV49239.1 |
| *Methanohalophilus euhalobius* | 401 | SNY18196.1 | SNX99917.1 |
| *Methanohalophilus sp.* | 401 | RSD36461.1 | RSD35796.1 |
| *Methanothrix thermoacetophila PT* | 400 | ABK14596.1 | ABK14717.1 |
| *Methanosarcina thermophila TM-1* | 400 | AKB14287.1 | AKB12510.1 |
| *Archaeoglobus veneficus SNP6* | 400 | AEA46764.1 | AEA46101.1 |
| *Methanohalophilus portucalensis* | 400 | ATU08507.1 | ATU08985.1 |
| *Methanohalophilus portucalensis FDF-1* | 400 | OJH49864.1 | OJH50401.1 |
| *Methanohalophilus portucalensis FDF-1* | 400 | RNI13322.1 | RNI11170.1 |
| *Methanohalophilus portucalensis FDF-1* | 400 | SMH33344.1 | SMH29520.1 |
| *Thermococcus sp._2319x1* | 400 | ALV62581.1 | ALV62414.1 |
| *Methanococcoides methylutens* | 400 | KGK99286.1 | KGK98787.1 |

| | | | |
|---|---|---|---|
| *Archaeoglobales archaeon* | 399 | RLI76561.1 | RLI75147.1 |
| *Methanohalophilus sp. DAL1* | 399 | OBZ34745.1 | OBZ35371.1 |
| *Methanohalophilus sp. T328-1* | 399 | KXS45754.1 | KXS46650.1 |
| *Methanosarcina sp. 795* | 399 | ALK05471.1 | ALK05944.1 |
| *Methanosarcina flavescens* | 399 | AYK14249.1 | AYK15939.1 |
| *Methanohalophilus sp. RSK* | 399 | RNI13775.1 | RNI12029.1 |
| *Methanosphaera cuniculi* | 399 | PAV06866.1 | PAV08199.1 |
| *Methanosphaera cuniculi* | 399 | PWL08622.1 | PWL08283.1 |
| *Methanohalophilus euhalobius* | 398 | PQV42226.1 | PQV43857.1 |
| *Methanohalophilus euhalobius* | 398 | RNI07908.1 | RNI11966.1 |
| *ANME-2 cluster archaeon* | 397 | PXF61384.1 | PXF60841.1 |
| *Methanococcoides methylutens MM1* | 395 | AKB84414.1 | AKB84801.1 |
| *Methanolobus vulcani* | 394 | SDF83415.1 | SDG03721.1 |
| *Methanosarcina sp. A14* | 393 | OEC90124.1 | OED09644.1 |
| *Thermococcus litoralis DSM 5473* | 393 | EHR78694.1 | EHR77470.1 |
| *Thermoprotei archaeon* | 392 | RLE88343.1 | RLE88080.1 |
| *Candidatus Verstraetearchaeota archaeon* | 391 | RLE52182.1 | RLE52385.1 |
| *Thermococcales archaeon_bit 391* | 391 | KUJ99511.1 | KUJ98857.1 |
| *Methanosaeta harundinacea 6Ac* | 390 | AET63426.1 | AET65143.1 |
| *Archaeoglobales archaeon* | 389 | RLI84427.1 | RLI86467.1 |
| *Methanocaldococcus sp. FS406-22* | 389 | ADC70293.1 | ADC69876.1 |
| *Pyrococcus horikoshii OT3* | 389 | BAA29387.1 | BAA30164.1 |
| *Methanosarcina sp. Kolksee* | 389 | AKB46910.1 | AKB47961.1 |
| *Methanosarcina horonobensis HB-1 = JCM 15518* | 389 | AKB77649.1 | AKB78609.1 |
| *Methanosarcina barkeri 3* | 389 | AKB82254.1 | AKB81067.1 |
| *Methanobacterium sp. 42_16* | 388 | KUK74549.1 | KUK74816.1 |
| *Methanobacterium formicicum* | 388 | AIS31832.1 | AIS33107.1 |
| *Methanosarcina mazei S-6* | 388 | AKB64182.1 | AKB64911.1 |
| *Archaeoglobus fulgidus DSM 4304* | 387 | AAB89032.1 | AAB90608.1 |
| *Methanocaldococcus jannaschii DSM 2661* | 387 | AAB99009.1 | AAB98949.1 |
| *Methanosphaera sp. WGK6* | 387 | OED30174.1 | OED30001.1 |
| *Methanonatronarchaeum thermophilum* | 387 | OUJ18801.1 | OUJ19107.1 |
| *Methanobrevibacter filiformis* | 387 | KZX10441.1 | KZX12494.1 |
| *Methanolobus psychrophilus R15* | 387 | AFV24914.1 | AFV22472.1 |
| *Methanococcoides burtonii DSM 6242* | 387 | ABE53049.1 | ABE51195.1 |
| *Methanobacterium sp. BAmetb5* | 386 | AXV38957.1 | AXV40205.1 |
| *Thermococcus onnurineus NA1* | 386 | ACJ16164.1 | ACJ17293.1 |
| *Methanosarcina vacuolata Z-761* | 386 | AKB43450.1 | AKB44460.1 |
| *Methanococcoides vulcani* | 386 | SES79485.1 | SES91445.1 |
| *Methanosarcina sp. Ant1* | 386 | OEU43252.1 | OEU42792.1 |
| *Methanobacterium sp. MZ-A1* | 385 | AUB58978.1 | AUB57684.1 |
| *Methanobacterium subterraneum* | 385 | AUB56150.1 | AUB55339.1 |
| *Methanotorris igneus Kol 5* | 385 | AEF96801.1 | AEF96386.1 |
| *Methanonatronarchaeia archaeon* | 385 | RZN61090.1 | RZN63254.1 |
| *Methanosarcina sp. 1.H.T.1A.1* | 384 | KKH99733.1 | KKH96291.1 |
| *Methanosarcina sp. 1.H.A.2.2* | 384 | KKH45833.1 | KKH45292.1 |
| *Methanosphaera sp. A6* | 384 | OEC85334.1 | OEC93525.1 |
| *Methanosphaera stadtmaniae DSM 3091* | 384 | ABC56899.1 | ABC57838.1 |
| *Thermoplasmata archaeon* | 384 | RLF67634.1 | RLF67227.1 |

| | | | |
|---|---|---|---|
| *Thermoprotei archaeon_bit383* | 383 | RLE60622.1 | RLE61176.1 |
| *Methanolobus tindarius DSM 2278* | 383 | ETA67130.1 | ETA66623.1 |
| *Methanosarcina sp. 2.H.A.1B.4* | 383 | KKG10365.1 | KKG08698.1 |
| *Methanolobus profundi* | 382 | SFM17242.1 | SFM32228.1 |
| *Thermococcus kodakarensis KOD1* | 382 | BAD85463.1 | BAD85937.1 |
| *Thermococcus chitonophagus* | 382 | ASJ17289.1 | ASJ16566.1 |
| *Thermococcus chitonophagus* | 382 | CUX77913.1 | CUX77525.1 |
| *Candidatus Syntrophoarchaeum caldarius* | 382 | OFV67164.1 | OFV68207.1 |
| *Methanosphaera sp. DEW79* | 382 | RAP48072.1 | RAP46203.1 |
| *Methanosarcina acetivorans C2A* | 382 | AAM06425.1 | AAM05817.1 |
| *Thermococcus profundus* | 381 | ASJ02920.1 | ASJ03378.1 |
| *Archaeoglobales archaeon ex4484_92* | 380 | OYT34858.1 | OYT38065.1 |
| *Palaeococcus pacificus DY20341* | 380 | AIF70188.1 | AIF68629.1 |
| *Methanosarcina lacustris Z-7289* | 380 | AKB75324.1 | AKB74393.1 |
| *Methanosarcina sp. WH1* | 380 | AKB21243.1 | AKB21505.1 |
| *Methanosarcina sp. WWM596* | 380 | AKB17905.1 | AKB18173.1 |
| *Pyrococcus kukulkanii* | 380 | AMM53559.1 | AMM54013.1 |
| *Thermoprotei archaeon* | 379 | RLE96880.1 | RLE98028.1 |
| *Methanocaldococcus vulcanius M7* | 379 | ACX72463.1 | ACX73357.1 |
| *Methanobacterium lacus* | 379 | ADZ09190.1 | ADZ08406.1 |
| *Methanosarcina sp. 2.H.T.1A.3* | 379 | KKG15605.1 | KKG16397.1 |
| *Methanosarcina sp. 2.H.T.1A.8* | 379 | KKG27506.1 | KKG27436.1 |
| *Methanosarcina sp. 2.H.T.1A.6* | 379 | KKG19533.1 | KKG21492.1 |
| *Methanosarcinales archaeon* | 379 | RLG35033.1 | RLG35959.1 |
| *Candidatus Methanomassiliicoccus intestinalis Issoire-Mx1* | 379 | AGN26388.1 | AGN26625.1 |
| *Methanocaldococcus bathoardescens* | 377 | AIJ05795.1 | AIJ06123.1 |
| *Candidatus Methanoperedens nitroreducens* | 377 | SNQ61043.1 | SNQ59840.1 |
| *Methanosarcina acetivorans* | 377 | RXA20151.1 | RXA16576.1 |
| *Methanocaldococcus infernus ME* | 376 | ADG13687.1 | ADG13320.1 |
| *Methanocaldococcus fervens AG86* | 376 | ACV24729.1 | ACV24654.1 |
| *Methanosarcina spelaei* | 375 | PAV11703.1 | PAV12371.1 |
| *Methanohalobium evestigatum Z-7303* | 374 | ADI73235.1 | ADI74739.1 |
| *Methanobrevibacter thaueri* | 374 | PWB87567.1 | PWB85415.1 |
| *Thermococcus guaymasensis DSM 11113* | 373 | AJC71925.1 | AJC71987.1 |
| *Methanosarcinales archaeon* | 372 | RLG31377.1 | RLG31003.1 |
| *Pyrococcus sp. ST04* | 372 | AFK22042.1 | AFK22376.1 |
| *Methanosarcina sp. MTP4* | 372 | AKB25727.1 | AKB25344.1 |
| *Methanosarcina siciliae T4/M* | 372 | AKB29385.1 | AKB28192.1 |
| *Methanomethylovorans hollandica DSM 15978* | 372 | AGB50046.1 | AGB49746.1 |
| *Thermococcus gammatolerans EJ3* | 371 | ACS33184.1 | ACS33090.1 |
| *Methanotorris formicicus Mc-S-70* | 371 | EHP89697.1 | EHP87372.1 |
| *Methanolobus sp. T82-4* | 371 | KXS44985.1 | KXS41885.1 |
| *Thermococcus gorgonarius* | 370 | ASJ00910.1 | ASJ01134.1 |
| *Methanobacterium subterraneum* | 369 | AUB59980.1 | AUB60817.1 |
| *Thermoprotei archaeon_Bitscore_369* | 369 | RLF05530.1 | RLF05913.1 |
| *Candidatus Methanoperedenaceae archaeon HGW-Methanoperedenaceae-1* | 369 | PKL53941.1 | PKL53787.1 |
| *Pyrococcus sp. NA2* | 369 | AEC51848.1 | AEC52416.1 |
| *Pyrococcus furiosus DSM 3638* | 369 | AAL80414.1 | AAL81220.1 |
| *Thermococcus pacificus* | 369 | ASJ07203.1 | ASJ06348.1 |

| | | | |
|---|---|---|---|
| *Thermococcus nautili* | 368 | AHL22193.1 | AHL23063.1 |
| *Methanosphaera sp. BMS* | 368 | AWX32291.1 | AWX33055.1 |
| *Thermococcus thioreducens* | 368 | ASJ12469.1 | ASJ11549.1 |
| *Methanobrevibacter millerae* | 368 | ALT68580.1 | ALT70001.1 |
| *Candidatus Methanoperedens nitroreducens* | 367 | KCZ72414.1 | KCZ72254.1 |
| *Methanosaeta sp. NSM2* | 366 | OYV12742.1 | OYV13106.1 |
| *Candidatus Methanoperedenaceae archaeon* | 366 | PWB56072.1 | PWB52493.1 |
| *Methanobacterium formicicum DSM 3637* | 365 | EKF86403.1 | EKF86854.1 |
| *Thermococcus barossii* | 365 | ASJ04600.1 | ASJ05698.1 |
| *Methanobrevibacter gottschalkii* | 365 | SEK47361.1 | SEK77896.1 |
| *Pyrococcus abyssi GE5* | 364 | CAB50557.1 | CAB49833.1 |
| *Methanosphaera sp. SHI1033* | 364 | RAP45300.1 | RAP43861.1 |
| *Thermococcus paralvinellae* | 363 | AHF80223.1 | AHF79437.1 |
| *Candidatus Syntrophoarchaeum sp. WYZ-LMO15* | 362 | RJS73567.1 | RJS73300.1 |
| *Thermococcus barophilus MP* | 362 | ADT83693.1 | ADT85127.1 |
| *Thermococcus sp. 40_45* | 362 | KUK29368.1 | KUK29194.1 |
| *Methanobrevibacter sp. A27* | 362 | OED00616.1 | OEC95823.1 |
| *Thermofilum sp. NZ13* | 361 | PLJ77248.1 | PLJ77807.1 |
| *Methanobrevibacter sp. YE315* | 361 | AMD17679.1 | AMD16632.1 |
| *Methanosarcinales archaeon* | 360 | RLG37468.1 | RLG39219.1 |
| *C.Methanoliparum thermophilum* | 360 | RZN64856.1 | RZN63930.1 |
| *Thermococcus peptonophilus* | 359 | AMQ19205.1 | AMQ18827.1 |
| *Thermofilum pendens Hrk 5* | 358 | ABL77734.1 | ABL77540.1 |
| *Thermococcus piezophilus* | 358 | ANF23623.1 | ANF22274.1 |
| *Thermococcus sp. EP1* | 358 | KPU63538.1 | KPU63428.1 |
| *Methanosarcinales archaeon* | 357 | RZB32910.1 | RZB28998.1 |
| *C.Verstraetearchaeota archaeon_bit357* | 357 | RLE54104.1 | RLE56160.1 |
| *Thermococcus sp. 5-4* | 357 | ASA77500.1 | ASA78610.1 |
| *Thermococci archaeon_B45_G15* | 357 | RLF85673.1 | RLF84003.1 |
| *Candidatus Methanoperedens sp. BLZ1* | 354 | KPQ44352.1 | KPQ45088.1 |
| *Pyrococcus yayanosii CH1* | 353 | AEH24496.1 | AEH24247.1 |
| *Methanosarcinales archaeon* | 353 | RZN33637.1 | RZN41520.1 |
| *Methanosarcinales archaeon* | 352 | RZN13757.1 | RZN15380.1 |
| *Candidatus Verstraetearchaeota archaeon* | 352 | RZN55384.1 | RZN57389.1 |
| *Candidatus Methanohalarchaeum thermophilum* | 352 | OKY78142.1 | OKY78566.1 |
| *Methanothrix soehngenii GP6* | 351 | AEB67882.1 | AEB68079.1 |
| *Thermococcus radiotolerans* | 350 | ASJ14045.1 | ASJ15348.1 |
| *Thermococcus sp. P6* | 350 | ASJ10686.1 | ASJ10274.1 |
| *Thermococcus sibiricus MM 739* | 350 | ACS90026.1 | ACS90178.1 |
| *Thermococcus siculi* | 350 | ASJ09210.1 | ASJ08328.1 |
| *Thermofilum carboxyditrophus 1505* | 348 | AJB42637.1 | AJB42444.1 |
| *Thermococcus cleftensis* | 348 | AFL96219.1 | AFL95079.1 |
| *Thermofilum uzonense* | 347 | AKG39233.1 | AKG38124.1 |
| *Candidatus Methanoperedens sp.* | 347 | TAN45321.1 | TAN42969.1 |
| *Thermococci archaeon_B89_G9* | 347 | RLF76783.1 | RLF76802.1 |
| *methanogenic archaeon mixed culture ISO4-G1* | 347 | AMK13600.1 | AMK13653.1 |
| *Candidatus Altiarchaeales archaeon* | 346 | RLI91478.1 | RLI90953.1 |
| *Candidatus Altiarchaeales archaeon* | 346 | RLI94130.1 | RLI95359.1 |
| *Thermococcus sp. 4557* | 346 | AEK72217.1 | AEK73129.1 |

| | | | |
|---|---|---|---|
| *Methanococcales archaeon HHB* | 345 | GBF36496.1 | GBF36521.1 |
| *Halalkalicoccus jeotgali B3* | 345 | ADJ16338.1 | ADJ13447.1 |
| *Candidatus Altiarchaeales archaeon* | 345 | RLI89155.1 | RLI89996.1 |
| *Methanosphaerula palustris E1-9c* | 344 | ACL17363.1 | ACL15545.1 |
| *Thermococci archaeon_B61_G1* | 343 | RLF94847.1 | RLF96291.1 |
| *Methanobrevibacter sp. 87.7* | 343 | OWT32853.1 | OWT33112.1 |
| *Thermococci archaeon_B88_G9_342^* | 342 | RLF89829.1 | RLF90615.1 |
| *Aciduliprofundum sp. MAR08-339* | 341 | AGB05360.1 | AGB04768.1 |
| *Crenarchaeota archaeon_bit341 13_1_40CM_2_52_14* | 341 | OLD34472.1 | OLD35717.1 |
| *Crenarchaeota archaeon 13_1_40CM_3_52_17* | 341 | OLD11591.1 | OLD11796.1 |
| *Thermococcus sp. AM4* | 339 | EEB74978.1 | EEB74787.1 |
| *Candidatus Diapherotrites archaeon* | 339 | MAG17978.1 | MAG18084.1 |
| *Thermococci archaeon_B48_G16* | 338 | RLF95608.1 | RLF94557.1 |
| *Halodesulfurarchaeum formicicum* | 337 | APE96138.1 | APE94570.1 |
| *Thermoprotei archaeon_Bitscore_336a* | 336 | RLF21101.1 | RLF23064.1 |
| *Thermoprotei archaeon_Bitscore_336b* | 336 | RLF10651.1 | RLF13159.1 |
| *Methanofollis sp. FWC-SCC2* | 334 | TAJ44321.1 | TAJ43460.1 |
| *Thermococci archaeon_B54_G1* | 334 | RLF95176.1 | RLF94338.1 |
| *Crenarchaeota archaeon 13_1_40CM_3_53_5* | 333 | OLD03773.1 | OLD02883.1 |
| *Thermoplasmata archaeon* | 333 | RLF64338.1 | RLF64905.1 |
| *Thermoprotei archaeon_B75_G16* | 331 | RLF07210.1 | RLF06241.1 |
| *Candidatus Altiarchaeales archaeon* | 331 | RLI90894.1 | RLI91905.1 |
| *Candidatus Methanosuratus sp.* | 331 | RWX73511.1 | RWX73251.1 |
| *Methanospirillum stamsii* | 328 | PWR75421.1 | PWR75023.1 |
| *Candidatus Diapherotrites archaeon CG08_land_8_20_14_0_20_34_12* | 328 | PIU21193.1 | PIU21525.1 |
| *Methanococcus maripaludis X1* | 327 | AEK19591.1 | AEK20495.1 |
| *Halobacteriales archaeon QS_1_68_20* | 326 | PSP78786.1 | PSP79117.1 |
| *Candidatus Diapherotrites archaeon CG11_big_fil_rev_8_21_14_0_20_37_9* | 326 | PIN84663.1 | PIN85743.1 |
| *Methanospirillum hungatei JF-1* | 325 | ABD42323.1 | ABD40364.1 |
| *Methanosarcina barkeri MS* | 325 | AKB53689.1 | AKB53987.1 |
| *Haladaptatus litoreus* | 325 | SIQ68254.1 | SIR55518.1 |
| *Methanothermococcus okinawensis IH1* | 324 | AEH06325.1 | AEH07341.1 |
| *Halobacteriales archaeon SW_9_67_24* | 324 | PSQ65474.1 | PSQ66138.1 |
| *Methanococcus maripaludis C7* | 323 | ABR66663.1 | ABR65786.1 |
| *Methanocalculus sp. MSAO_Arc2* | 323 | RQD81624.1 | RQD83950.1 |
| *Methanobacterium sp. Maddingley MBC34* | 322 | EKQ54166.1 | EKQ51213.1 |
| *Candidatus Altiarchaeales archaeon* | 321 | RLI85163.1 | RLI86951.1 |
| *Crenarchaeota archaeon 13_1_20CM_2_53_14* | 321 | OLE58720.1 | OLE58596.1 |
| *Candidatus Altiarchaeales archaeon IMC4* | 321 | ODS42941.1 | ODS42669.1 |
| *Halorientalis regularis* | 321 | SDF42512.1 | SDG26691.1 |
| *Thermoprotei archaeon_Bitscore_320* | 320 | RLE77360.1 | RLE73932.1 |
| *Natronomonas pharaonis DSM 2160* | 319 | CAI48235.1 | CAI48396.1 |
| *Candidatus Aenigmarchaeota archaeon* | 318 | RLI96491.1 | RLI98574.1 |
| *Natronomonas moolapensis 8.8.11* | 318 | CCQ37866.1 | CCQ37696.1 |
| *Candidatus Altiarchaeales archaeon ex4484_2* | 317 | OYT53902.1 | OYT54696.1 |
| *Halobacterium salinarum NRC-1* | 315 | AAG20602.1 | AAG20324.1 |
| *Candidatus Bathyarchaeota archaeon RBG_16_48_13* | 315 | OGD46437.1 | OGD46372.1 |
| *Halobacteriales archaeon QH_9_66_26* | 315 | PSP72024.1 | PSP71108.1 |
| *Halarchaeum sp. CBA1220* | 313 | RNH90812.1 | RNH91068.1 |

| | | | |
|---|---|---|---|
| *Methanomicrobiales archaeon HGW-Methanomicrobiales-2* | 313 | PKL61908.1 | PKL62643.1 |
| *Halorientalis sp. F13-25* | 313 | RXK47292.1 | RXK50172.1 |
| *Halobacterium jilantaiense* | 313 | SEW13405.1 | SEW19187.1 |
| *Methanoregula boonei 6A8* | 312 | ABS56238.1 | ABS56783.1 |
| *Halococcus thailandensis JCM 13552* | 312 | EMA51912.1 | EMA53101.1 |
| *Methanophagales archaeon* | 311 | PXF53390.1 | PXF52694.1 |
| *Methanoculleus sp. MAB1* | 311 | CVK32226.1 | CVK31469.1 |
| *Halorientalis persicus* | 311 | SEO34681.1 | SEN47689.1 |
| *Methanocalculus sp. MSAO_Arc1* | 311 | RQD79351.1 | RQD81728.1 |
